# Nature exposure induces hypoalgesia by acting on nociception-related neural processing

**DOI:** 10.1101/2024.04.29.591600

**Authors:** Maximilian O. Steininger, Mathew P. White, Lukas Lengersdorff, Lei Zhang, Alexander J. Smalley, Simone Kühn, Claus Lamm

**Affiliations:** Social, Cognitive and Affective Neuroscience Unit, Department of Cognition, Emotion, and Methods in Psychology, Faculty of Psychology, University of Vienna, Vienna, Austria; Cognitive Science Hub, University of Vienna, Vienna, Austria; European Centre for Environment and Human Health, University of Exeter, UK; Environment and Climate Research Hub, University of Vienna, Vienna, Austria; Centre for Human Brain Health, School of Psychology, University of Birmingham, Birmingham, UK; Institute for Mental Health, School of Psychology, University of Birmingham, Birmingham, UK; Lise Meitner Group for Environmental Neuroscience, Max Planck Institute for Human Development, Berlin, Germany; Department of Psychiatry, University Medical Center Hamburg-Eppendorf, Hamburg, Germany

**Author notes:** **Corresponding author:** Claus Lamm.

**Keywords:** nature exposure, nature benefits, pain, neuroimaging, biological markers

## Abstract

Nature exposure has numerous psychological benefits, and previous findings suggest that exposure to nature reduces self-reported acute pain. Given the multi-faceted and subjective quality of pain and methodological limitations of prior research, it is unclear whether the evidence indicates genuine hypoalgesia or results from domain-general effects and subjective reporting biases. This preregistered functional neuroimaging study aimed to identify how nature exposure modulates nociception-related and domain-general brain responses to acute pain. We compared the self-reported and neural responses of healthy neurotypical participants (N = 49) receiving painful electrical shocks while exposed to virtual nature or to closely matched urban and indoor control settings. Replicating existing behavioral evidence, pain was reported to be lower during exposure to the natural compared to the urban or indoor control settings. Crucially, machine-learning-based multi-voxel signatures of pain demonstrated that this subjective hypoalgesia was associated with reductions in nociception-related rather than domain-general cognitive-emotional neural pain processing. Preregistered region-of-interest analyses corroborated these results, highlighting reduced activation of areas connected to lower-level somatosensory aspects of pain processing (such as the thalamus, secondary somatosensory cortex, and posterior insula). These findings demonstrate that nature exposure results in genuine hypoalgesia and that neural changes in lower-level nociceptive pain processing predominantly underpin this effect. This advances our understanding of how nature may be used as a non-pharmacological pain treatment. That this hypoalgesia was achieved with brief and easy-to-administer virtual nature exposure has important practical implications and opens novel avenues for research on the precise mechanisms by which nature impacts our mind and brain.

## Introduction

Natural settings such as parks, woodlands, coastlines, and natural elements, including plants, sunsets, and natural soundscapes, can protect and promote a range of health and well-being outcomes (1–3). People who live in greener neighborhoods tend to react less strongly to stressors (4) and have better mental health in the long term (5), regular nature visitors report fewer negative and higher positive emotional states (6), and even short experimental nature exposures can positively impact subjective and neural indicators of well-being (7). Theories connecting nature and health underscore various aspects that render certain natural environments particularly salutary. While stress recovery theory (SRT) proposes that the presence of natural, non-threatening content elicits positive affective responses and aids recovery from stress (8), attention restoration theory (ART) puts a stronger emphasis on nature’s ability to replenish voluntary attentional resources (9). According to ART, nature encompasses numerous elements that captivate human attention in a unique and effortless way. While differing in focus, both theories highlight nature’s capacity for human health, an assumption that has been substantiated by a multitude of evidence.

Of particular relevance to this study, natural settings may even have the potential to reduce acute pain (10–12). Forty years ago, Ulrich (1984) showed that patients recovering from surgery were given fewer analgesics to manage pain, had more positive healthcare provider notes, and left the hospital earlier when having a window view of trees compared to a brick wall (10). Similar results have subsequently been reported using various forms of nature exposure during diverse pain-related settings (e.g., invasive medical procedures such as dental treatments or bronchoscopy; 11, 12). However, the evidence to date has several limitations.

For instance, due to a lack of proper experimental controls previous work has been unable to fully assess whether it is nature specifically that reduces pain. Most studies have either not compared nature exposure to an alternative stimulation or used control conditions that were not carefully matched on key aspects such as low- or high-level visual features or subjective beauty (13, 14). For example, nature is often juxtaposed with aesthetically unpleasing or stressful settings, such as unappealing and busy urban environments. It thus remains unclear whether natural scenes reduce pain or if the alternative environments exacerbate it through their negative characteristics (8). To conclusively assess this possibility, sophisticated and carefully controlled experimental designs are required, ensuring that natural and alternative stimulations are closely matched on relevant key features.

Furthermore, most prior research has relied on self-report measures of pain, which, whilst important, are limited in two central regards. First, self-reports make it challenging to capture the multi-faceted quality of pain. Pain entails several components, ranging from lower-level sensory aspects, such as nociception and its neural processing, to higher-level components, involving affective, cognitive, and motivational processes and their associated neural responses (15). The sensory aspects predominantly reflect people’s ability to identify from where in the body a painful stimulus originated, how intense it is, and what type of pain is perceived. The cognitive-affective and motivational aspects entail feelings of unpleasantness towards the stimulus and the inclination to engage in protective behavior, as well as pain-related affect regulation. Although separate ratings of pain intensity and unpleasantness aiming to disentangle these aspects on a subjective level can be obtained experimentally (16), such self-reports are susceptible to various confounding influences (17). Second, affective, cognitive, and motivational processes associated with pain also play a role in other types of subjective experiences and thus may not entirely reflect pain-specific but rather domain-general processing (18). We cannot exclude that previous findings were primarily driven by the effects of nature on such domain-general processes and, therefore, lack specificity for pain. Moreover, self-report is limited by individual constraints in self-perception and meta-cognition, and beliefs about how nature exposure will influence one’s pain sensitivity and other types of experimental demand effects may have unintentionally influenced prior findings (19).

Neuroimaging techniques have thus been suggested as a possible way to complement self-report and facilitate a systems-level approach to the brain bases of pain. Indeed, experiencing pain involves numerous interconnected brain structures, and particular brain regions may be associated with distinct pain components (20). For example, while the posterior insula (pINS) and the secondary somatosensory cortex (S2) are linked to early lower-level nociception-related processing, higher-level components incorporating emotional or motivational aspects are associated with regions such as the anterior midcingulate (aMCC) and the prefrontal (PFC) cortex (20, 21). Evaluating the activation of these areas during acute pain could yield more refined and less subjective assessments of the various processes underpinning the multifaceted quality of pain and help to disentangle if lower- or higher-level processes are impacted.

In this respect, recent advancements in pain research are of particular value. For example, machine learning approaches and multivariate brain patterns have been applied to neuroimaging data to identify and differentiate between various aspects of pain with even higher precision and validity when compared to the analysis of single isolated brain regions (22).

Specifically, two prominent multivoxel patterns, the neurologic pain signature (NPS; 22), and the stimulus intensity independent pain signature-1, (SIIPS1; 23) have been developed to investigate and differentiate between lower-level and higher-level pain-related processing, respectively. The NPS tracks the intensity of a painful stimulus and involves brain regions that receive nociceptive afferents (24), thus tracing processes closely connected to nociception and lower-level sensations. The SIIPS1 has been developed to assess pain-related brain activity beyond nociception and, therefore, captures aspects such as motivational value and emotional or cognitive context (23). Importantly, the NPS has been shown to predict pain individually with high sensitivity and specificity, allowing the disambiguation from non-specific processes such as negative emotion or cognitive appraisal. To date, however, these recent methodological developments and neuroscientific insights have not been exploited to understand better the neural processes and mechanisms by which nature exposure might lead to the reduction of painful experiences. Besides advancing our basic knowledge, such research may have considerable importance for efforts to complement pharmaceutical treatment approaches, with their well-documented negative side effects and addictive properties (25).

To address these research gaps, we conducted a preregistered repeat-crossover functional magnetic resonance imaging (fMRI) experiment (preregistration: osf.io/t8dqu). In the fMRI scanner, healthy human participants were exposed to carefully matched virtual natural and urban scenes, as well as an indoor setting control condition, while experiencing electric shocks that induced individually calibrated acute transient pain (Figure 1). Combining multivoxel brain signature approaches (both NPS and SIIPS1) with analyses of distinct pain-responsive brain areas allowed us to explore the impact of nature stimuli (vs. urban and indoor controls) on different aspects of the pain-processing hierarchy.

**Figure 1.**
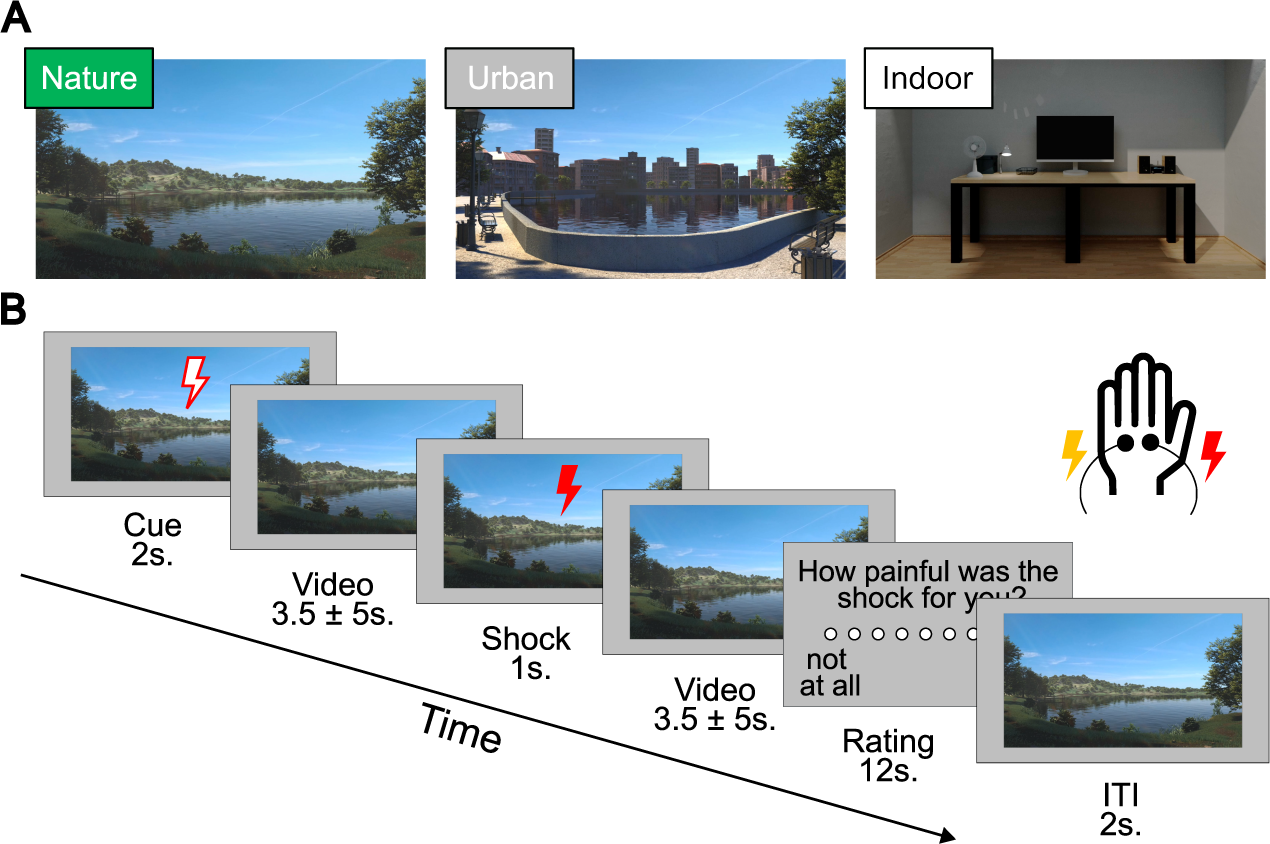
Stimuli and trial structure of the experiment. (*A*) Stimuli depicting a natural, an urban, and an indoor environment. A matching soundscape accompanied each visual stimulus. The three pain runs had a total duration of 9 minutes each, during which one environment was accompanied by 16 painful and 16 nonpainful shocks. All participants were exposed to all environments (in counterbalanced order). (*B*) Structure and timeline of an example trial. First, a cue indicating the intensity of the next shock (red = painful, yellow = not painful) was presented for 2,000 ms. Second, a variable interval of 3,500 ± 1,500 ms was shown. Third, a cue indicating the intensity of the shock was presented for 1,000 ms, accompanied by an electrical shock with a duration of 500 ms. Fourth, a variable interval of 3,500 ± 1,500 ms followed. Fifth, after each third trial, participants rated the shock’s intensity and unpleasantness at 6,000 ms each. Sixth, each trial ended with an intertrial interval (ITI) presented for 2,000ms. The environmental stimulus was presented simultaneously except for the rating phase during each trial. Electrical painful and non-painful shocks were administered to the dorsum of the left hand with a separate electrode. <insert page break here>

Based on previous research, yet using a sophisticated experimental design with highly controlled experimental stimuli, we hypothesized that exposure to nature compared to urban or indoor control settings would reduce self-reported pain. For the neuroimaging data, with which we aimed to significantly extend previous behavioral research, we predicted that pain-related neural activity would be reduced by exposure to nature compared to the control conditions. Both hypotheses were preregistered. While we expected reductions in brain responses associated with lower-level nociception-related or higher-level pain-related emotional-cognitive processes, the lack of prior neuroimaging research precluded specific predictions about which of the two processes would be impacted preferentially.

## Results

### Nature stimuli reduce self-reported pain

We used immediate self-report ratings of experienced pain intensity and unpleasantness to study participants’ subjective pain response. With the intensity ratings, we intended to capture the sensory-discriminate, and thus nociception-related, aspects of pain, while the unpleasantness ratings aimed to measure higher-level cognitive-emotional and motivational features (16, 21). Participants were carefully instructed to discriminate both aspects and rated each separately on a scale from zero (“not at all painful/unpleasant”) to eight (“very painful/unpleasant”; see *Experimental Procedures*). Statistical inferences of the self-report data were based on linear mixed modeling (LMM; see *Methods and Material* and *Supporting Information*).

Supporting our preregistered hypothesis, we found a significant main effect of *environment* (nature, urban, or indoor) on the immediate ratings [i.e., pooled intensity and unpleasantness ratings, F_(2,48)_ = 12.48, p < 0.001)]. Planned pairwise contrasts revealed that self-reported pain was lower in the nature vs. urban [b = −0.54, SE = 0.12, t = −4.46, p < 0.001 one-tailed, d_rm_ = −0.59] and indoor condition [b = −0.48, SE = 0.1, t = −4.14, p < 0.001 one-tailed, d_rm_ = −0.52], with urban and indoor conditions not differing [b = 0.06, SE = 0.12, t = 0.52, p = 0.69, d_rm_ = 0.01]. We found a significant interaction effect of *environment*rating type* [F_(2,81.14)_ = 9.19, p < 0.001)]. Investigations of the beta parameters and planned pairwise contrasts suggested that this interaction reflected that the magnitude, but not the overall pattern regarding how the three environment conditions affected the two types of ratings, differed. As displayed in Figures 2A-B, the differences between nature and the other two conditions were larger for the unpleasantness than for the intensity ratings, with effect sizes representing medium-to-high and small magnitudes, respectively. Specifically, planned pairwise contrasts revealed a significant difference in intensity ratings between nature vs. urban [b = −0.25, SE = 0.12, t = −2.14, p = 0.018 one-tailed, d_rm_ = −0.29] and nature vs. indoor [b = −0.29, SE = 0.11, t = −2.67, p = 0.005 one-tailed, d_rm_ = −0.33] but not for urban vs. indoor [b = −0.04, SE = 0.12, t = −0.38, p = 0.71, d_rm_ = −0.05]. Similarly, the unpleasantness ratings showed a significant difference comparing nature vs. urban [b = −0.83, SE = 0.16, t = −5.23, p < 0.001 one-tailed, d_rm_ = −0.86] and nature vs. indoor [b = −0.66, SE = 0.15, t = −4.35, p < 0.001 one-tailed, d_rm_ = −0.69], but again not when comparing urban vs. indoor [b = 0.17, SE = 0.15, t = 1.12, p = 0.27, d_rm_ = 0.17]. In addition to the immediate intensity and unpleasantness ratings, participants were asked to assess retrospectively (directly after concluding a complete pain block, i.e., exposure to an environment coupled with painful shocks) to what extent viewing the respective environments helped them distract themselves from or better tolerate the shocks. These ratings revealed a significantly higher level of distraction from and tolerance of the shocks for the nature condition compared to both the urban and indoor conditions. No difference between urban vs. indoor was found (see *Supporting Information* for statistics).

**Figure 2.**
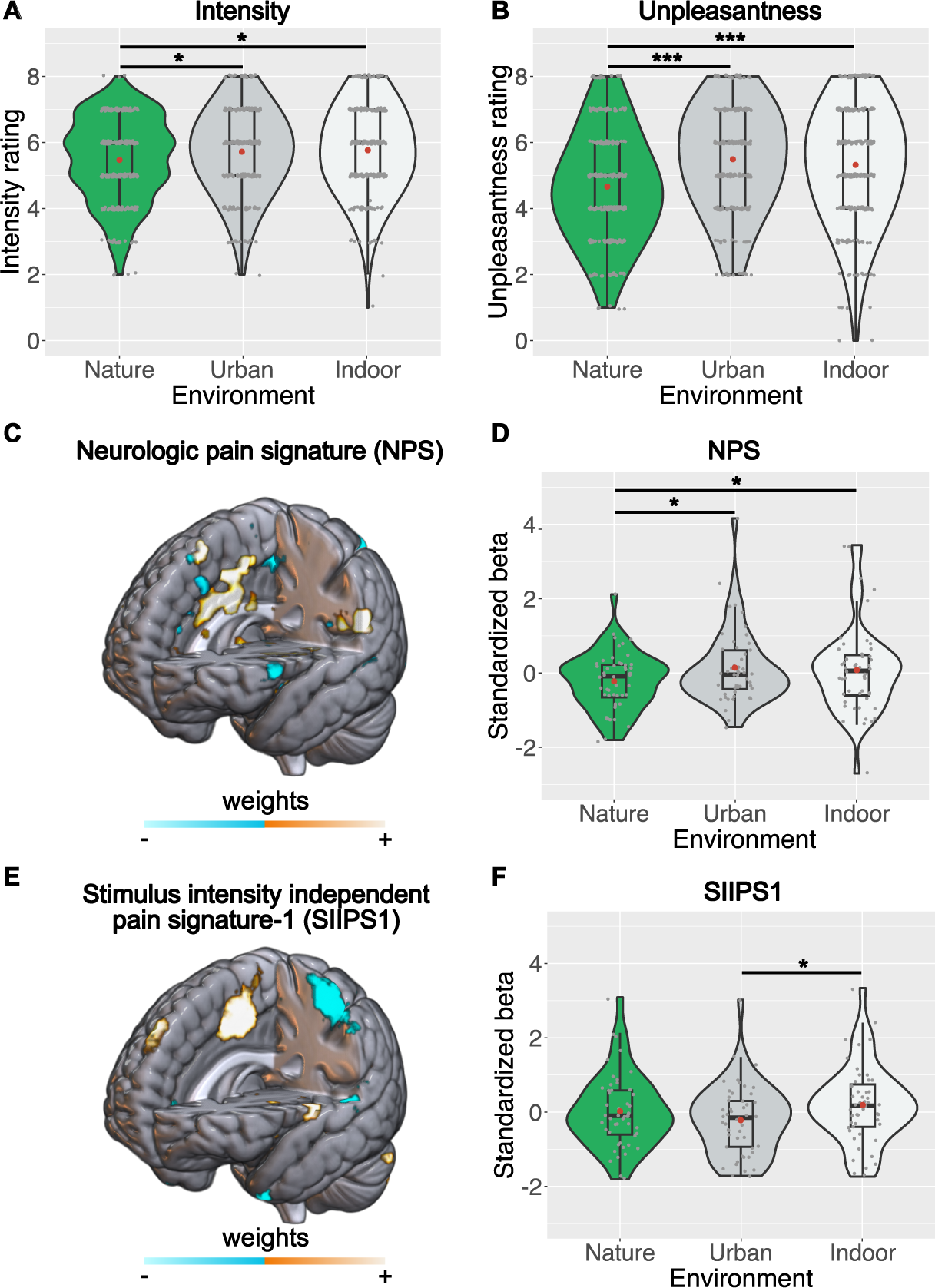
Violin plots depicting (*A*) intensity and (*B*) unpleasantness ratings of painful shocks and the overall lower-level nociceptive (*D*) and higher-level cognitive-emotional (*F*) neural response to pain as indicated by the neurologic pain signature (NPS, *C*) and the stimulus intensity independent pain signature-1 (SIIPS1, *E*) for each environment. Both brain maps show the signatures’ weights (positive = orange, negative = blue). For display purposes, the map of the SIIPS1 shows weights that exceed a predefined threshold (false discovery rate of q < 0.05). Intensity and unpleasantness ratings were given on a scale from 0 (“not at all painful/unpleasant”) to 8 (“very painful/unpleasant”). NPS and SIIPS1 responses are plotted as standardized beta values. Grey and red dots in violin plots represent single values and mean scores, respectively. * < .05, ** < .01, *** < .001, mark significant planned pairwise comparisons derived from the linear mixed models.

These results confirm our preregistered hypotheses and go beyond prior findings of self-reported pain reduction. They indicate that the change in pain is specific to a decrease in the natural setting, rather than an increase in the alternative settings. Furthermore, within the typical constraints of self-reports, the immediate ratings suggest that both sensory-discriminative (indicated by intensity) and affective-motivational (indicated by unpleasantness) processing was impacted similarly, but that the latter showed a more pronounced effect. The retrospective ratings provide the additional insight that participants perceived nature stimuli as helping them with pain tolerance via attention distraction.

### Nature stimuli reduce nociception-related neural responses to pain

We first clearly confirmed that the pain paradigm effectively engaged brain signatures and regional responses classically associated with neural pain processing (see *Supporting Information* showing significant NPS and SIIPS1 as well as region of interest responses for pain vs. no pain across all three conditions). We then assessed the main hypothesis that exposure to nature vs. control stimuli differently affects multivoxel signatures of lower-level nociception-related or higher-level cognitive-emotional responses to pain. To this end, we first computed the NPS and the SIIPS1 in each environmental condition and then compared them using LMM with the signatures per condition as the dependent variable. We found no significant result for the main effect of *environment* [F_(2,48)_ = 1.25, p = 0.296)], but, importantly, a significant interaction effect of *environment*signature* [F_(2,96)_ = 6.04, p = 0.003], indicating that the environments impacted the NPS and SIIPS1 differently. Specifically, planned pairwise contrasts revealed significant decreases in the NPS response during nature compared to urban [b = −0.37, SE = 0.16, t = −2.27, p = 0.013 one-tailed, d_rm_ = −.41] and indoor environments [b = −0.30, SE = 0.18, t = −1.68, p = .049 one-tailed, d_rm_ = −.30], with low to moderate effect sizes. There was no significant effect when comparing urban vs. indoor environments [b = 0.07, SE = 0.16, t = 0.39, p = .69, d_rm_ = .06]. For the SIIPS1, no significant effects for the nature vs. urban or indoor comparison were found (p = .93 and p = .17, both one-tailed; see *Supporting Information*), but a significant difference of urban vs. indoor [b = −0.41, SE = 0.16, t = 2.49, p = 0.014, d_rm_ = −.41] (see Figure 2D and 2F).

The signature-based analyses provided important insights into how the three different environments affected comprehensive neural activation patterns related to pain. Inspired by recent multiverse approaches of neuroimaging data (26) aiming to identify converging evidence across complementary analysis approaches, we had planned and preregistered additional analyses of specific regions of interest (ROIs) and how their activation was affected by the three environments. Selection of the ROIs was theory-based, covering key areas of three circuits involved in the processing and modulation of pain (see *Materials and Methods*) identified in an influential framework for pain research (21). The first circuit represents the ascending pathway and includes the primary somatosensory cortex (S1) and the thalamus. The two other circuits represent descending modulatory systems engaged by psychological pain alterations. One circuit encompasses the superior parietal lobe (SPL), secondary somatosensory cortex (S2), posterior insula (pINS), and amygdala and is associated with attentional modulations of pain. The other circuit covers the anterior insula (aINS), anterior midcingulate cortex (aMCC), medial prefrontal cortex (mPFC) and periaqueductal gray (PAG) and is engaged when emotions alter pain.

Analyzing each of the ROIs separately using a LMM revealed the following significant results for the main effect of *environment*: Thalamus [F_(2,48)_ = 5.53, p = 0.007], S2 [F_(2,48)_ = 5.16, p = 0.009], pINS [F_(2,48)_ = 9.28, p = 0.0003], and a trend for the amygdala [F_(2,48)_ = 2.68, p = 0.078]. Planned pairwise contrasts revealed a significant difference when comparing nature vs. urban in the thalamus [b = −0.28, SE = 0.11, t = −2.25, p = 0.014 one-tailed, d_rm_ = −0.39], S2 [b = −0.50, SE = 0.18, t = −2.78, p = 0.008 one-tailed, d_rm_ = −0.47], pINS [b = −0.96, SE = 0.23, t = −4.25, p < 0.001 one-tailed, d_rm_ = −0.78] and the amygdala [b = −0.17, SE = 0.09, t = −1.89, p = 0.042 one-tailed, d_rm_ = −0.34]. Comparing nature vs. indoor revealed a significant difference in the thalamus [b = −0.38, SE = 0.12, t = −3.18, p = 0.003 one-tailed, d_rm_ = −0.48], S2 [b = −0.39, SE = 0.17, t = − 2.33, p = 0.038 one-tailed, d_rm_ = −0.36], pINS [b = −0.38, SE = 0.19, t = −1.92, p = 0.038 one-tailed, d_rm_ = −0.31], and the amygdala [b = −0.12, SE = 0.06, t = −1.80, p = 0.038 one-tailed, d_rm_ = −0.28]. None of the remaining ROIs showed significant differences for the main effect of environment (all p-values > .125, see *Supporting Information*). Calculating planned pairwise contrasts between urban vs. indoor for the ROIs reported above also revealed no significant differences (see *Supporting Information*).

In summary, the multivoxel and region of interest analyses converge in showing that pain responses when exposed to nature as compared to urban or indoor stimuli are associated with a decrease in neural processes related to lower-level nociception-related features (NPS, thalamus), as well as in regions of descending modulatory circuitry associated with attentional alterations of pain that also encode sensory-discriminative aspects (S2, pINS).

## Discussion

This preregistered neuroimaging study investigated whether exposure to nature vs. urban or indoor control stimuli mitigates subjective and neural responses to acute pain. Using carefully selected and designed control stimuli and leveraging neuroimaging techniques allowed us to address two potential major confounds of previous findings. First, that the less appealing and more aversive quality of the contrasting stimuli rather than the positive qualities of nature explained the observed changes in pain. Second, that constraints associated with subjective pain measures, such as reporting biases or experimental demand effects, confounded earlier results. Furthermore, drawing upon a comprehensive preregistered analysis approach of our fMRI data enabled us to specifically identify the neural responses to pain that were positively affected by nature exposure.

Following this approach, we demonstrate that natural settings, compared to matched urban or indoor scenes, induce genuine hypoalgesia, that the effects are positive consequences of the nature stimuli rather than being caused by the aversiveness of the standard ‘urban’ control stimuli, and that this effect can be attributed to changes at sensory and nociception-related lower levels of the processing hierarchy. More specifically, nature exposure was associated with a reduced response in a highly precise and sensitive neurological signature of pain (the NPS) linked to nociception-related brain processes (22). Complementary univariate analyses showed lowered pain-related activation in areas receiving nociceptive afferents (thalamus, S2, pINS), providing converging evidence that nature exerted its effects predominantly on areas associated with lower-level sensory pain components. Moreover, the stimulus-intensity independent pain signature-1 (SIIPS1), used to capture higher-level pain-related processes, was not differentially affected by the nature stimuli, further supporting that nociception-related rather than cognitive-emotional aspects underpinned the subjective hypoalgesia. Importantly, these novel neural findings were corroborated by reduced self-reported pain, replicating past research (11, 12, 27).

Regarding our first preregistered hypotheses, we replicate and crucially extend the specificity of previous findings by demonstrating that comparing virtual nature to a matched urban and an additional neutral indoor scene leads to consistent patterns of reduced self-reported pain. Including two control conditions and showing that pain ratings were lower in the nature setting (but similar in the urban and indoor scene), we find that alterations in pain are attributable to a decrease in the nature condition rather than an increase in the urban one - a confound that seems particularly plausible as most urban environments are associated with increased stress levels (8). Importantly, unlike most past work, we used pre-tested and published stimuli of closely matched natural and urban settings (see *Materials and Methods*) that were both rated as comparably beautiful (28). Specifically, the urban stimuli contained many appealing and attractive elements from the nature scene, reducing the possibility that any differences would result from merely creating a spatially unmatched, noxious, and aesthetically unpleasing urban setting (13, 14). Furthermore, using both immediate and retrospective pain ratings, we show that this change in self-report is consistent across different indicators of subjective pain.

The consistency of immediate and retrospective ratings is important because it convergently validates the experimental effects and reveals important intuitions and introspective insights by the participants into how the three environments may have influenced their pain experience and its regulation. Specifically, that participants thought the nature scenes helped to distract them from the pain, and in this way, to tolerate the shocks better is an aspect that converges with attention-related neural processes as a possible mechanism of reduced nociceptive pain that we will discuss further below. However, the immediate ratings of intensity and unpleasantness also reveal why it is important to complement self-report using neural data (19). Indeed, while both types of ratings showed a decrease in the nature setting, effect sizes were higher for unpleasantness than intensity ratings. This suggests that nature influenced the affective-motivational rather than the sensory-discriminative components of pain (21), which is not corroborated by our neural findings. A possible explanation of this discrepancy is that the self-report may reflect participants’ assumptions about how the different environments will affect their experiences. In particular, given that the nature stimuli elicit stronger positive affect (28), participants may have assumed and reported diminished negative affective pain. Since subjective ratings are the result of an intricate interplay between various mechanisms (including nociception, emotion, or cognition), using such ratings alone would make it difficult to conclude which specific aspect of pain processing was impacted (17).

Leveraging highly sensitive neural indicators of specific pain components helped us overcome this limitation. Using these neural indicators demonstrates that the decreased subjective reports of pain are associated with reduced neural responses in lower-level nociceptive pain, as indicated by a selective effect on the neurologic pain signature (NPS). This is a key finding, as the NPS entails several regions that receive nociceptive afferents and shows high pain specificity. This is indicated by a lack of responsiveness to a range of experiences that are related but not specific to pain, such as cognitive appraisal and aversive affect (19, 22). There is thus broad consensus that experimental manipulations that result in changes of this signature indicate genuinely pain-related, and in particular nociception-related, brain states (though see 18 for a critical account). Importantly, the NPS effects are also, like the self-report effects, specifically related to the nature stimuli and not confounded by increases in pain processing due to inappropriately matched urban and indoor control stimuli, respectively.

Beyond demonstrating pain specificity, comparing the NPS with another pain signature, the SIIPS1, revealed that nature acted predominantly on nociception-related rather than domain-general aspects of pain. The SIIPS1 has been developed to capture pain-related processes as well, but in contrast to the NPS characterizes domain-general cognitive and affective aspects of pain beyond nociception-related and somatosensory processing (23). Pain regulation or valuation are two examples of such aspects, which are linked to ventral and dorsal prefrontal cortex activity and thus to higher-level associative brain areas farther removed from the direct somatosensory inputs (29). Therefore, it is another key insight that this signature, and how it tracked the acute pain we exposed our participants to, was not affected differentially by the nature vs. control stimuli. It should be noted, though, that we only preregistered the investigation regarding the NPS but not the SIIPS1 since we had originally planned to disentangle which pain components are predominantly affected using pooled ROIs activity. This decision was adopted later, but before looking at the data, because a direct comparison between signatures upon further reflection seemed more parsimonious and valid (see *Supporting Information*).

As our study was the first to use these neuroimaging approaches to investigate the underlying neural processes of nature-based hypoalgesia, we regard the selective effects on the NPS as requiring confirmation by further research. However, the complementary analyses of individual ROIs strengthen the signature-based findings that nature exposure acts on lower-rather than higher-level pain processing. Of note, these ROI analyses were planned with two rationales in mind. First, in the spirit of multiverse analyses (26), they aimed to analyze our data in different ways and render our conceptual conclusions more convincing if convergent evidence was revealed. Second, they allowed us to tap into distinct pathways connected to pain and its neural representation. Compared to the more data-driven brain signatures, these neural pathways are based on long-standing theoretical accounts grounded in pain physiology and clinical practice (15, 21). Drawing upon these accounts, we find decreased activation during the nature condition in the ascending pathway (thalamus) receiving direct input from nociceptors and a descending modulatory circuit involving areas associated with sensory-discriminative processing (e.g., S2, pINS). In contrast, brain regions related to a circuit underlying higher-level emotional modulations of pain (e.g., aMCC, mPFC) showed no difference between environments. This is important because it enables us to disentangle the underlying mechanisms, relate the findings to influential accounts of the benefits of nature from environmental psychology, and put them into perspective relative to other non-pharmacological interventions.

For instance, in the most extensive single neuroimaging study of placebo effects to date, it was suggested that placebo manipulations do not impact nociception-related (NPS), but instead domain-general cognitive-emotional aspects (SIIPS1) of pain (30). This is in direct contrast to our findings and suggests that nature-related pain reductions are likely not based on belief processes such as the ones investigated by placebo research. Instead, pain relief through nature exposure seems to be more related to changes in sensory circuitries and attentional processes connected to the engagement of these circuits. Similar results have been found among participants engaged in attention-based mindfulness practices (31), where training participants in mindfulness practices over eight weeks was associated with changes in lower-level nociception-related (NPS) but not higher-level cognitive-emotional (SIIPS1) responses to pain. The authors interpreted this reduced NPS response as changes in attentional mechanisms that gate lower-level nociceptive signals.

Regarding nature’s potential to alleviate pain, the interpretation that reduced NPS activity is indicative of altered attentional processing is particularly intriguing. In the field of environmental psychology, changed attentional processing is indeed one of the proposed key mechanisms linking nature exposure to health (9). Attention restoration theory (ART) suggests that natural stimuli can “restore” depleted attentional capacities. The reasoning behind this argument is that nature possesses many features that are “softly fascinating” to humans and engage us in a distracting but not overly demanding manner. In the context of pain, this implies that natural features have the potential to capture attention in unique ways, thereby diverting it away from the painful sensation more effectively than other environments. In conjunction with findings from neuroscientific pain research, the observed reduction in nociception-related responses substantiates this interpretation in two ways.

First, neuroscientific accounts of pain propose that different modulatory neural systems are engaged when pain is altered by emotional or attentional processes (21). For instance, previous studies have shown that if attention is diverted from a painful stimulus, this is visible in changed responses in areas related to sensory-discriminative processing (32–34; for a critical account see 35). According to these frameworks, attentional modulations of pain are characterized by pathways involving projections from the superior parietal lobe to the insula, S2, and amygdala (21). We observed effects (or trends towards them, for the amygdala) for most of these areas when comparing nature to urban or indoor stimuli. Second, asking participants if exposure to the respective environment helped to distract themselves from pain better revealed effect sizes in the medium to high range when comparing nature to urban (d_rm_ = .66) or indoor settings (d_rm_ = 1.04) while comparing urban and indoor stimuli (d_rm_ = 0.34) showed only a small effect (see *Supporting Information*). Together, these theoretical accounts and our findings render it plausible that the effects on nociceptive signaling and its cortical representations are linked to attention-related processes. However, it should be noted that an attention regulation mechanism and the precise pattern of results were not specifically preregistered. The postulated interaction between attention- and nociception-related processes thus needs confirmation and extension by future research, which should focus on identifying how exactly attention-related brain areas act as regulators of the nociceptive inputs.

Besides this proposition for future work, our findings open several other exciting research avenues. First, participants in our study were not exposed to real-world environments but to virtual stimuli. While this approach allowed us to maximize experimental control, whether the results are generalizable to real-world contexts remains to be tested. That our findings are based on virtual stimuli is a major strength, though. It suggests that nature-based therapies do not necessarily require real-world exposure, but that stimuli acting as proxies for such environments might suffice. This is a particularly promising aspect as it suggests a broad range of use cases that can be employed cost-efficiently in a wide range of interventions.

Second, more granularity is required to thoroughly assess which specific elements of nature are relevant in driving the observed hypoalgesia. The literature on the benefits of nature suggests that certain perceptual features make natural settings particularly fascinating (9, 13). These features might exhibit a notably engaging effect, thus leading to a stronger diversion from pain. Furthermore, the complex cognitive and emotional reactions, such as feelings of awe and nostalgia, towards these features might be essential (36). Further work is thus needed to explore which specific sensory elements make natural environments particularly effective in alleviating pain.

Third, while harnessing neuroimaging enabled us to interrogate the effects of natural settings on pain processing with unprecedented specificity, some accounts challenge the notion that neuroimaging indicators can entirely dissociate pain from other phenomena (18). To further increase the specificity of the evidence that nature impacts nociceptive pain, future studies may use additional measures to expand on the specific components and processes nature affects. In this respect, it would be intriguing to test patients suffering from congenital insensitivity to pain, a condition characterized by absent nociceptive processing. If the subjective hypoalgesia is truly grounded in changes in nociception-related processing, these patients should, compared to our neurotypical sample, not be impacted by the natural settings.

Finally, the current work focused on the modulation of acute pain. Given the severe impact chronic pain has on patients and our society and the potential risks associated with its pharmacological treatment, nature exposure represents an interesting complementary pain management strategy. While the current study provides first evidence as to which underlying processes are altered in the processing of acute pain, chronic pain is characterized by complex and multifaceted changes in psychological and neural processing (37) that only partially converge with those during acute pain. Thus, future research should investigate if and by which mechanisms exposure to nature might help to alleviate chronic pain conditions.

In conclusion, our results show that simple and brief exposure to nature reduces self-reported and specific neural responses to acute pain and is linked to lower-level pain-specific nociception-related processing. In contrast to other non-pharmacological interventions, which usually involve complex deceptions through placebo induction procedures or week-long training of cognitive coping strategies, the nature stimuli used here potentially provide an easily accessible alternative or at least complimentary intervention in clinical practice. Incorporating natural elements into healthcare design has the potential to reduce pain-associated complaints and constraints with relatively low effort. This is important and promising from a clinical-applied perspective: it suggests that employing natural stimuli could be a cost-effective and easily implementable intervention in pain treatment and related contexts to promote health and well-being.

## Materials and Methods

### Participants

The study was conducted according to the seventh revision of the Declaration of Helsinki (2013) and approved by the Ethics Committee of the University of Vienna (EK-Nr. 00729). A total of 53 healthy right-handed human participants fulfilling standard inclusion criteria for neuroimaging studies of pain participated. Based on an a-priori power analysis, a sample size of 48 participants was preregistered (see *Supporting Information*). Four participants had to be excluded due to technical problems with the pain stimulator and the scanner, leading to a final sample including 24 female and 25 male participants (Age ± SD = 25.24 ± 2.79, range = 20-35). All participants received a reimbursement of €30.

### Experimental Procedures

Upon arrival, participants were instructed about the study procedure, gave written informed consent, and completed a pain calibration task. Afterward, they entered the MRI scanner and were alternately exposed to blocks of virtual stimuli, each depicting a different environment, directly followed by blocks showing the same environment accompanied by electrical shocks (from here on referred to as “video” and “pain” blocks, respectively). This design enabled participants to familiarize themselves with each respective environment before its presentation alongside the pain stimuli.

To deliver an engaging and immersive experience each stimulus was created by a dedicated professional graphic designer and depicted a virtual environment accompanied by a matching soundscape. Three different environments were presented in counterbalanced order, showing a natural, an urban, or an indoor setting (see Figure 1A). The natural and urban environments were closely matched regarding low-level (e.g., color and spatial properties) and high-level (e.g., scenic structure, complexity, openness) visual features (28). Specifically, the natural setting was created first and included a large central lake (with observable wind ripples), trees by the side of the lake (with rustling leaves), and an animation showing the shifting position of the sun and cloud movements. The urban condition was constructed by adding human-made elements to this basic scene, including buildings on the far side of the lake, a paved path, a short wall, and benches on the nearside of the lake. The resulting urban scene, containing many of the originally attractive natural elements, was still rated as relatively beautiful (28). Both scenes were accompanied by soundscapes created based on recommendations of previous works investigating acoustic experiences in different environments (38). The nature scene included the sounds of rippling water, gentle wind, native birds, and insects, while the urban scene included the sounds of different vehicles and construction works. For both environments, careful consideration was given to selecting and adjusting all sounds based on factors such as the nativeness of species, typical local traffic noises (e.g., emergency vehicle horns), or the time of day. The indoor setting depicted a desk with office supplies, a fan, and a computer. It was accompanied by the sounds of a computer and a fan. The soundscapes of all environments were normalized regarding their average loudness by matching the root-mean-square amplitude. To further increase the level of immersion, we instructed participants to imagine themselves being present in the specific environment by reading through a short script preceding each block. The script was based on previous nature-based guided imagery interventions (39).

During pain blocks, participants re-watched the same environment but additionally received electrical shocks. Thirty-two electrical shocks (16 painful and 16 non-painful) were administered per block. To ensure comparable pain intensities across participants, the stimuli were calibrated according to an established procedure (40, 41). Painful shocks were calibrated to represent a “very painful, but bearable” (6), and non-painful shocks to represent a “perceptible, but non-painful” (1) sensation on a scale from 0 (“not perceptible”) to 8 (“unbearable pain”). We administered the shocks using a Digitimer DS5 Isolated Bipolar Constant Current Stimulator (Digitimer Ltd, Clinical & Biomedical Research Instruments). Two electrodes, one for painful and one for non-painful shocks, were attached to the dorsum of the left hand. Mean shock intensities were 0.61 mA (SD = 0.42) and 0.19 mA (SD = 0.09) for painful and non-painful trials, respectively, which is comparable to previous studies in our laboratory following a similar protocol (40, 42). Each pain block presented the painful and non-painful trials in the same pseudorandomized order. Pseudorandomization was employed to ensure that the co-occurrence of painful shocks and specific auditory and visual elements of the environments were kept constant across participants and conditions. In line with previous uses of the pain paradigm (40, 42), every trial started with a colored visual cue displayed for 2,000 ms that indicated the next shock’s intensity (painful = red, non-painful = yellow). After a variable pause where the cue disappeared (jittered with 3,500 ± 1,500 ms), another visual cue was presented for 1.000 ms with the electrical stimulus being administered for 500 ms simultaneously. The second visual cue matched the first cue in shape and size but had a colored filling. Next, the cue and shock disappeared for a variable duration (jittered with 3,500 ± 1,500 ms). An additional intertrial interval of 2,000 ms separated all trials (Figure 1B). Twelve of the 32 trials (six painful and six non-painful) were succeeded by two ratings to indicate the perceived intensity (“How painful was the shock for you?”) or unpleasantness (“How unpleasant was the shock for you?”) of the last administered shock on a scale ranging from zero (“not at all”) to eight (“very”). Notably, the visual cues for each trial were superimposed on the virtual scene, which continuously played in the background to maximize the immersion into the environment. The visual and accompanying audio stimuli were presented on an MRI-compatible 32-inch display (Full HD 1920×1080 PPI resolution; BOLDscreen 32 LCD, Cambridge Research System, Cambridge, UK) viewed at 26° x 15° visual angle, and Sensimetrics earphones (model S14; Sensimetrics Corporation, Gloucester, MA, USA), respectively. All stimuli and ratings were presented using MATLAB R2021a (Mathworks, 2021) and Psychophysics Toolbox Version 3 (43).

### fMRI Acquisition, Preprocessing, and Analysis

fMRI data were acquired with a 3 Tesla Siemens Magnetom Skyra MRI scanner (Siemens Medical, Erlangen, Germany). The scanner was equipped with a 32-channel head coil. Each run acquired a separate functional volume for one of the three pain blocks using the following parameters: Repetition time (TR) = 800 ms, echo time (TE) = 34 ms, flip angle = 50°, field of view (FOV) = 138 mm, multi-band acceleration factor = 4, interleaved multi-slice mode, interleaved acquisition, matrix size = 96 × 96, voxel size = 2.2 × 2.2 × 3.5 mm^3^, 36 axial slices of the whole brain, and slice thickness = 3.85 mm. We used a magnetization-prepared rapid acquisition gradient echo sequence with the following parameters to obtain the structural image at the end of each scanning session: TR = 2.300 ms, TE = 2.29 ms, flip angle = 8°, FOV = 240 mm, ascending acquisition, single shot multi-slice mode, 256 sagittal slices, voxel size = 0.94 × 0.935 × 0.935 mm3, slice thickness = 0.935 mm.

Preprocessing of the fMRI data was performed using SPM12 (Wellcome Trust Centre for Neuroimaging, www.fil.ion.ucl.ac.uk/spm) running on MATLAB 2021a (Mathworks, 2021), including the following steps: realignment and unwarping using participant-specific field maps, slice-time correction with the center slice as reference, coregistration of functional and structural images, segmentation into three tissue types (gray matter, white matter, cerebrospinal fluid), spatial normalization to Montreal Neurological Institute space using Diffeomorphic Anatomical Registration Through Exponentiated Lie Algebra (DARTEL), and spatial smoothing with a 6-mm full-width at half maximum 3D Gaussian Kernel. The first-level analyses followed a general linear model (GLM) approach. A design matrix was specified in which the painful and non-painful trials were modeled as experimental regressors per environment (i.e., run). Furthermore, six nuisance regressors from the realignment step accounting for movement-induced noise were added per environment. The experimental regressors were time-locked to the onset of each shock and convolved using SPM12’s standard hemodynamic response function in an event-related fashion.

To ascertain that our pain paradigm, as expected and extensively demonstrated in prior work (20, 22, 23, 44), robustly activated single-region and multivariate signature responses to pain, we first performed an analysis that was orthogonal to our main hypotheses. This analysis revealed conclusive evidence that our pain task evoked neural activity in pain-related brain regions, the NPS and SIIPS1, and all preregistered ROIs except the left S1 (ipsilateral to the stimulated hand; see *Supporting Information*). Therefore, we proceeded to test our main hypotheses on whether these neural responses to pain are reduced by exposure to nature. To this end, one contrast image was created comparing pain > no-pain trials for each environment. First, we investigated whether the overall lower-level nociception-related and higher-level cognitive-emotional neural response to pain differed for each environment by applying the NPS and the SIIPS1 to our first-level GLM beta maps (22). This was done using scripts created by the developers of these patterns (22, 23), which were made available to us after personal enquiry. We calculated the dot product of the contrast image and the pattern map of the NPS and SIIPS1, resulting in two scalar values for each participant and environment. The NPS and SIIPS1 represent multivoxel patterns within and across pain-related brain regions that track lower-level or higher-level pain processing, respectively (22, 23). Second, we performed ROI analyses to test our hypotheses using a different methodological approach and to further differentiate if the alterations in pain are predominantly found in areas associated with lower-level or higher-level pain processing. We created the following preregistered set of sphere-based ROIs (center [± x, y, z]; sphere size): amygdala (±20, −12, −10]; 10mm), anterior midcingulate cortex (aMCC; [-2, 23, 40], 10mm), anterior insula (aINS; [±33, 18, 6]; 10mm), posterior insula (pINS; [±44, −15, 4]; 10mm), medial prefrontal cortex (mPFC; [7, 44, 19]; 10mm), primary somatosensory cortex (S1; [±39, −30, 51]; 10mm), secondary somatosensory cortex (S2; [±39, −15, 18]; 10mm), periaqueductal gray (PAG; [0, −32, −10]; 6mm), superior parietal lobe (SPL; [±18, −50, 70]; 10mm), and thalamus ([±12, −18, 3]; 6mm). Each ROI’s center coordinate and sphere size were based on previous meta-analytic findings and pain studies from our lab experimentally inducing acute pain using similar methods (40, 42, 45). For each ROI, we only included voxels that showed a significant response to painful vs. non-painful stimuli in the pain>no-pain contrast across environments. Then, we extracted the mean percent signal change per participant for the pain>no-pain first-level contrasts for each individual environment using the MarsBar toolbox (46).

### Statistical Analysis

To test our main hypothesis, which was that exposure to nature stimuli reduces self-report and neural responses to pain, we ran several LMMs using the lmer function of the lme4 package in R (47). We preregistered the majority of the models (osf.io/t8dqu) and specified each of them using maximal random effects structures (48). For the immediate self-reports on pain, we specified the intensity and unpleasantness ratings of the painful shocks as the dependent variable to be predicted by the fixed effect of environment (nature as the reference), rating content (intensity as the reference), and their interaction (with random slopes and intercepts for environment, rating content and their interaction by participant). For the neural signatures, we used the NPS and SIIPS1 as the dependent variable to be predicted by the fixed effect of environment (nature as a reference), signature (NPS as reference), and their interaction (with random slopes and intercepts for environment and signature by participant). For ROIs in one hemisphere, we used the ROI response as the dependent variable to be predicted by the fixed effect of environment (nature as a reference, with random intercepts for participants). For ROIs with spheres in both hemispheres, we used the ROI responses of both hemispheres as the dependent variable to be predicted by the fixed effect of environment (nature as a reference), hemisphere (left as reference), and their interaction (with random slopes and intercepts for environment and hemisphere by participant). For each LMM, we report significance testing for the main effects of environment and interaction effects of interest, followed by planned pairwise comparisons. The p-values of the pairwise comparisons from the ROI analysis were Bonferroni-Holm corrected (separated by the different descending modulatory (attention vs. emotion) and ascending pain circuits; all reported p-values represent adjusted values). For each pairwise comparison, we computed the repeated standardized mean difference (d_rm_) as an effect size using the means and standard deviations of each environment (49). An exemplary model syntax, using the response in the S2 as a dependent variable, looked like this:

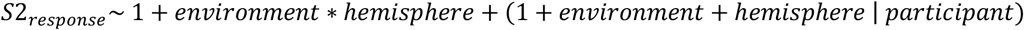

Details regarding all models (e.g., formulae, model fit, random effects variance and correlation, etc.) and deviations from the preregistration are reported in the *Supporting Information*.

## Acknowledgments

This research was funded by the Austrian Science Fund (FWF: W1262-B29; M3166). MW’s time on this project was supported by the EU’s Horizon Europe research and innovation programme under grant agreement No. 101081420 (RESONATE). Participants were, in part, recruited through the Vienna CogSciHub: Study Participant Platform (SPP), based on the Hamburg Registration and Organization Online Tool (hroot; Bock et al., 2014).

## Author Contributions

M.O.S. conceptualization, methodology, software, validation, formal analysis, investigation, data curation, writing – original draft, writing – review & editing, visualization, project administration. M.P.W. conceptualization, methodology, writing – original draft, writing – review & editing, supervision. L.L. formal analysis, resources, writing – review & editing. L.Z. methodology, formal analysis, resources, writing – review & editing. A.J.S. resources, writing – review & editing. S.K. writing – review & editing. C.L. conceptualization, methodology, resources, writing - original draft, writing - review & editing, supervision, funding acquisition.

## Data availability

Behavioral data, region of interest, and multivariate signature data extracted from the fMRI signal course, as well as unthresholded statistical maps for the pain>no-pain contrast in each environment are asccessible at https://osf.io/t8dqu/.

## References

1. S. J. Geiger, et al., Coastal proximity and visits are associated with better health but may not buffer health inequalities. Commun Earth Environ 4, 166 (2023).

2. T. Hartig, R. Mitchell, S. de Vries, H. Frumkin, Nature and health. Annu Rev Public Health 35, 207–228 (2014).

3. C. Twohig-Bennett, A. Jones, The health benefits of the great outdoors: A systematic review and meta-analysis of greenspace exposure and health outcomes. Environ Res 166, 628–637 (2018).

4. F. Lederbogen, et al., City living and urban upbringing affect neural social stress processing in humans. Nature 474, 498–501 (2011).

5. R. S. Geary, et al., Ambient greenness, access to local green spaces, and subsequent mental health: A 10-year longitudinal dynamic panel study of 2.3 million adults in Wales. Lancet Planet Health 7, e809–e818 (2023).

6. M. P. White, et al., Associations between green/blue spaces and mental health across 18 countries. Sci Rep 11, 8903 (2021).

7. G. N. Bratman, J. P. Hamilton, K. S. Hahn, G. C. Daily, J. J. Gross, Nature experience reduces rumination and subgenual prefrontal cortex activation. Proc Natl Acad Sci USA 112, 8567–8572 (2015).

8. R. S. Ulrich, et al., Stress recovery during exposure to natural and urban environments. J Environ Psychol 11, 201–230 (1991).

9. S. Kaplan, The restorative benefits of nature: Toward an integrative framework. J Environ Psychol 15, 169–182 (1995).

10. R. Ulrich, View through a window may influence recovery from surgery. Science 224, 420–421 (1984).

11. G. B. Diette, N. Lechtzin, E. Haponik, A. Devrotes, H. R. Rubin, Distraction therapy with nature sights and sounds reduces pain during flexible bronchoscopy: A complementary approach to routine analgesia. Chest 123, 941–948 (2003).

12. K. Tanja-Dijkstra, et al., The soothing sea: A virtual coastal walk can reduce experienced and recollected pain. Environ Behav 50, 599–625 (2018).

13. K. E. Schertz, M. G. Berman, Understanding nature and its cognitive benefits. Curr Dir Psychol Sci 28, 496–502 (2019).

14. K. L. Meidenbauer, et al., The affective benefits of nature exposure: What’s nature got to do with it? J Environ Psychol 72, 101498 (2020).

15. N. Mercer Lindsay, C. Chen, G. Gilam, S. Mackey, G. Scherrer, Brain circuits for pain and its treatment. Sci Transl Med 13, eabj7360 (2021).

16. D. D. Price, Psychological and neural mechanisms of the affective dimension of pain. Science 288, 1769–1772 (2000).

17. C.-W. Woo, T. D. Wager, What reliability can and cannot tell us about pain report and pain neuroimaging. Pain 157, 511–513 (2016).

18. I.-S. Lee, E. A. Necka, L. Y. Atlas, Distinguishing pain from nociception, salience, and arousal: How autonomic nervous system activity can improve neuroimaging tests of specificity. Neuroimage 204, 116254 (2020).

19. M. C. Reddan, T. D. Wager, Modeling pain using fMRI: From regions to biomarkers. Neurosci Bull 34, 208–215 (2018).

20. A. V. Apkarian, M. C. Bushnell, R.-D. Treede, J.-K. Zubieta, Human brain mechanisms of pain perception and regulation in health and disease. Eur J Pain 9, 463–463 (2005).

21. M. C. Bushnell, M. Čeko, L. A. Low, Cognitive and emotional control of pain and its disruption in chronic pain. Nat Rev Neurosci 14, 502–511 (2013).

22. T. D. Wager, et al., An fMRI-based neurologic signature of physical pain. N Engl J Med 368, 1388–1397 (2013).

23. C.-W. Woo, et al., Quantifying cerebral contributions to pain beyond nociception. Nat Commun 8, 14211 (2017).

24. X. Han, et al., Effect sizes and test-retest reliability of the fMRI-based neurologic pain signature. Neuroimage 247, 118844 (2022).

25. W. M. Compton, N. D. Volkow, Abuse of prescription drugs and the risk of addiction. Drug Alcohol Depend 83, S4–S7 (2006).

26. R. Botvinik-Nezer, et al., Variability in the analysis of a single neuroimaging dataset by many teams. Nature 582, 84–88 (2020).

27. J. Stanhope, M. F. Breed, P. Weinstein, Exposure to greenspaces could reduce the high global burden of pain. Environ Res 187, 109641 (2020).

28. A. J. Smalley, M. P. White, Beyond blue-sky thinking: Diurnal patterns and ephemeral meteorological phenomena impact appraisals of beauty, awe, and value in urban and natural landscapes. J Environ Psychol 86, 101955 (2023).

29. V. Neugebauer, V. Galhardo, S. Maione, S. C. Mackey, Forebrain pain mechanisms. Brain Res Rev 60, 226–242 (2009).

30. R. Botvinik-Nezer, et al., Placebo treatment affects brain systems related to affective and cognitive processes, but not nociceptive pain. bioRxiv [preprint] (2023). https://www.biorxiv.org/content/10.1101/2023.09.21.558825 (accessed 15 April 2024).

31. J. Wielgosz, et al., Neural signatures of pain modulation in short-term and long-term mindfulness training: A randomized active-control trial. Am J Psychiatry 179, 758–767 (2022).

32. M. Ploner, M. C. Lee, K. Wiech, U. Bingel, I. Tracey, Flexible cerebral connectivity patterns subserve contextual modulations of pain. Cereb Cortex 21, 719–726 (2011).

33. C. Villemure, M. C. Bushnell, Mood influences supraspinal pain processing separately from attention. J Neurosci 29, 705–715 (2009).

34. P. Rainville, Brain mechanisms of pain affect and pain modulation. Curr Opin Neurobiol 12, 195–204 (2002).

35. D. M. Torta, V. Legrain, A. Mouraux, E. Valentini, Attention to pain! A neurocognitive perspective on attentional modulation of pain in neuroimaging studies. Cortex 89, 120–134 (2017).

36. A. J. Smalley, et al., Soundscapes, music, and memories: Exploring the factors that influence emotional responses to virtual nature content. J Environ Psychol 89, 102060 (2023).

37. L. E. Simons, I. Elman, D. Borsook, Psychological processing in chronic pain: A neural systems approach. Neurosci Biobehav Rev 39, 61–78 (2014).

38. E. Ratcliffe, Sound and soundscape in restorative natural environments: A narrative literature review. Front Psychol 12, 570563 (2021).

39. N. Weinstein, A. K. Przybylski, R. M. Ryan, Can nature make us more caring? Effects of immersion in nature on intrinsic aspirations and generosity. Pers Soc Psychol Bull 35, 1315–1329 (2009).

40. M. Rütgen, et al., Placebo analgesia and its opioidergic regulation suggest that empathy for pain is grounded in self pain. Proc Natl Acad Sci USA 112, E5638–E5646 (2015).

41. S. C. C. Chan, C. C. H. Chan, A. S. K. Kwan, K. Ting, T. Chui, Orienting attention modulates pain perception: An ERP study. PLoS One 7, e40215 (2012).

42. H. Hartmann, M. Rütgen, F. Riva, C. Lamm, Another’s pain in my brain: No evidence that placebo analgesia affects the sensory-discriminative component in empathy for pain. Neuroimage 224, 117397 (2021).

43. D. H. Brainard, The Psychophysics Toolbox. Spatial Vis 10, 433–436 (1997).

44. A. Xu, et al., Convergent neural representations of experimentally-induced acute pain in healthy volunteers: A large-scale fMRI meta-analysis. Neurosci Biobehav Rev 112, 300–323 (2020).

45. C. Lamm, J. Decety, T. Singer, Meta-analytic evidence for common and distinct neural networks associated with directly experienced pain and empathy for pain. Neuroimage 54, 2492–2502 (2011).

46. M. Brett, J.-L. Anton, R. Valabregue, J.-B. Poline, Region of interest analysis using an SPM toolbox. Neuroimage 16, 1140–1141, (2002).

47. D. Bates, M. Mächler, B. Bolker, S. Walker, Fitting linear mixed-effects models using lme4. J Stat Soft 67(2015).

48. D. J. Barr, R. Levy, C. Scheepers, H. J. Tily, Random effects structure for confirmatory hypothesis testing: Keep it maximal. J Mem Lang 68, 255–278 (2013).

49. D. Lakens, Calculating and reporting effect sizes to facilitate cumulative science: A practical primer for t-tests and ANOVAs. Front Psychol 4 (2013).

