## Supplement for "Nature exposure induces hypoalgesia by acting on nociception-related neural processing"

### Materials and Methods

**Inclusion Criteria for Participants.** Participants fulfilling the following criteria were included in the study: right-handedness, no past or present neurological, psychiatric, or chronic disorder, no past or current disorder affecting the perception of pain, no intake of psychopharmacological medication (including pain medication), no substance abuse (alcohol, drugs) within the last three months, no past or present enrollment in studies including pharmaceutics, medicine or psychology, no pregnancy, age between 18 and 35 years. Furthermore, standard criteria for being able to take part in a functional magnetic resonance imaging (fMRI) experiment had to be fulfilled by the participants.

**Power Analysis.** We conducted an a-priori power analysis, which yielded a planned sample size of N = 48 participants. This sample size was based on a power analysis conducted for a repeated measures ANOVA model using G*Power 3.1 (1). Although the preregistered and reported statistical models in this work are linear mixed models (LMM; 2, 3), the project was powered for repeated measures ANOVAs, which would have served as a fallback in case of convergence issues in the LMMs. Since, compared to repeated measures ANOVA, LMMs in most situations have a higher power, the estimated sample size from the analyses can be seen as a conservative estimate for the targeted LMMs. The power analysis was based on previous studies investigating the effect of different environmental stimuli on pain perception (4–7). The average of reported effect sizes comparing differences between nature stimuli to a complete absence of stimulation while experiencing pain was used (Cohen`s d = .65). Studies directly comparing natural to urban environments in the domain of pain research revealed an effect size of similar magnitude (Cohen's d = .71) as the average reported for the remaining studies (see reference 12 in main text). Using a type-I error probability of $\alpha = 0.05$, a power of $1-\beta = 0.8$, an $\varepsilon = 0.34$ for non-sphericity corrections, and conservative estimates for repeated measures correlation of r = 0.4 between consecutive pain measurements (8–10), we estimated a targeted sample size of N = 41 participants. Due to published studies possibly overestimating effect sizes due to publication/survivorship bias, we decided to use a larger sample size of N = 48.

**Deviations from and clarifications regarding the preregistration.** We deviated from the preregistration regarding the specification of our LMMs in the following points.

First, most of our preregistered LMMs were specified without adhering to the principle of using maximal random effect structures. However, it is recommended to define maximal random effect structures when using LMM for confirmatory hypothesis testing, as the model outputs tend to generalize best under these circumstances (see reference 48 in main text). Since confirmatory hypothesis testing was the intended aim of the current study, we decided to keep the random effect's structure maximal when possible and justified by the design. This modification had the added benefit of overcoming convergence issues in the models we encountered using the initially specified random effect structures from the preregistration (which included only random intercept per participant). Model formulae for each conducted LMM can be found in the respective tables of this document (Table S1-S2 and S5-S16).

Second, concerning the LMM of the immediate pain ratings (i.e., intensity and unpleasantness of individual shocks), we preregistered a model that included ratings of both painful and non-painful shocks. However, including non-painful shocks in the model led to convergence issues and violations of the model assumptions, possibly due to limited variation in the ratings of non-painful shocks (which were, as intended, predominantly rated as 0 or 1). Thus, we adapted the model and specified it excluding non-painful shocks. We believe that excluding the non-painful stimuli from the analysis does not compromise the relevance of our replication of previous findings in self-report. First, our study's main aim was to replicate how exposure to nature affects the processing of painful stimuli. Second, previous studies investigate how nature exposure impacts the processing of painful stimuli and do not examine nor report how these relate to sensory stimuli below the pain threshold.

Third, regarding the LMM of our neuroimaging data investigating overall neural responses to pain, we deviated in terms of the dependent variable used. To examine whether nature exposure acts on neural indicators of lower-level or higher-level components of pain, we preregistered an analysis using the pooled activation of various ROIs as a dependent variable.

The main aim of this analysis was to investigate whether the nature condition impacts the overall neural response to pain clustered into two main components (i.e., sensory-discriminative vs. affective-motivational). For this analysis, we specified a LMM in which all ROI responses related to sensory-discriminative (i.e., S1, S2, pINS, SPL) or affective-motivational (i.e., aMCC, aINS, mPFC, dlPFC, PAG) components were used as the dependent variable simultaneously. We included environment (nature as a reference), component (sensory-discriminative as a reference), and their interaction as fixed effects. Using this specification, we expected a significant interaction effect between environment and component, indicating that our nature condition would differentially impact the pooled activity of single ROIs clustered into sensory-discriminative vs. affective-motivational components. However, considering the feedback of colleagues, revisiting the literature on the neuroimaging of pain, and upon further reflection we decided to use multivoxel pattern responses instead of pooled ROI activity to disentangle nature's effect on lower-level or higher-level pain processing. To this end, we used the signal of two established multivoxel pain signatures, the neurologic pain signature (NPS) and the stimulus intensity independent pain signature-1 (SIIPS1), as a dependent variable (see main manuscript). In comparison to using single ROI responses, these multivariate patterns differentiate between the two components of pain with increasing precision and validity and operationalize them with high levels of specificity. Although we did preregister to compare the impact of our environments on the NPS we did not preregister using SIIPS1. We believe that using a more precise and valid operationalization of our targeted constructs increased the validity and parsimony of our findings. Nevertheless, we also calculated the originally planned LMM using the pooled activity of the single ROIs as a dependent variable and the fixed effects as specified above. As for all other models, we set the random effects structure maximal with random slopes and intercepts for environment and component by participant. Using this model, we corroborated the findings of the main manuscript by showing that nature, compared to urban or indoor scenes, also reduces pain when using a different indicator of lower-level sensory-discriminative pain processing. For details, see the results section of this document.

Finally, we would like to clarify a potentially ambiguous wording used in our preregistration. In the preregistration, we specified the following hypothesis: Viewing virtual nature stimuli leads to a downregulation of pain primarily via activity changes in either sensory-discriminative (i.e., S1, S2, SPL, pINS) or affective-motivational (i.e., aMCC, mPFC, dlPFC, aINS, PAG) pain components. We specified that our hypothesis regarding nature's impact on which of these two neural modulatory systems are affected is non-directional in the sense that a lack of prior studies precluded predicting with confidence whether lower-level or higher-level pain processes (or both) would be reduced. The non-directionality of this hypothesis is thus related to the interaction effect of the independent variables (i.e., environment*component). This interaction effect was also tested two-sidedly in our analysis. However, the pairwise planned contrasts that investigated whether each component separately showed a reduced response (as indicated by the wording "downregulation of pain" in the preregistration) were directional, which is why they were assessed with one-sided testing.

### Results

**Self-reported pain.** The following section reports all details regarding the LMM of the self-reported pain data for immediate (i.e., intensity and unpleasantness) as well as recollected (i.e., distraction and tolerance) ratings (see Table S1 and S2).

Regarding the immediate ratings, we calculated a LMM specifying the ratings of the painful shocks as the dependent variable (i.e., intensity and unpleasantness) to be predicted by the fixed effect of environment (nature as the reference), rating content (intensity as the reference), and their interaction (with random slopes and intercepts for environment, rating content and their interaction by participant). We observed significant main effects of environment [$F_{(2,48)} = 12.48$, $p < 0.001$)], rating content [$F_{(1,48)} = 17.51$, $p < 0.001$)], and an interaction of environment*content [$F_{(2,81.14)} = 9.19$, $p < 0.001$)]. The main effect of environment and its interaction with rating content, as well as the planned pairwise contrasts, are presented in the main manuscript. The planned pairwise contrasts for the main effect of rating content indicated that intensity (M = 5.65, SE = 0.12) was rated higher than unpleasantness (M = 5.16, SE = 0.16), irrespective of the environment shown [b = 0.49, SE = 0.12, t = 4.18, p < 0.001].

Regarding the retrospective ratings, participants were asked to give an overall assessment of how much viewing each respective environment helped them to tolerate better and distract themselves from the pain using two separate questions. This assessment was given at the end of each pain block, i.e., after watching one environment while receiving the electrical shocks. Participants answered the following two questions using a scale from one ("not at all") to five ("very"): "Sitting in [nature / the city / the room] distracted me from the electrical shocks." and "Sitting in [nature / the city / the room] helped me to better tolerate the electrical shocks.". We used this wording as a way to explicitly prompt participants to envision themselves in the respective environment before starting each pain block using a structured script. To analyze the data, we used a LMM (see Table S2). We specified the recollected ratings (i.e., distraction and tolerance) as the dependent variable to be predicted by the fixed effect of environment (nature as the reference), rating content (distraction as the reference), and their interaction (with random slopes and intercepts for environment and rating content by participant). The significance tests for the main effects and their interaction revealed that there was no significant overall effect of environment [$F_{(2,106)} = 0.95$, $p = 0.388$)], but a significant main effect of rating content [$F_{(1,48)} = 8.42$, $p = 0.005$)], and it's interaction with environment [$F_{(2,96)} = 3.38$, $p = 0.038$)]. Planned pairwise contrasts revealed that ratings of distraction (M = 2.44, SE = 0.09) compared to tolerance (M = 2.21, SE = 0.08) were rated higher, irrespective of the environment shown [b = 0.23, SE = 0.08, t = 2.90, p = 0.005, $d_{rm} = 0.36$]. Planned pairwise contrasts of the interaction effect revealed that comparing nature vs. urban [b = 0.69, SE = 0.20, t = 3.40, p = 0.014, $d_{rm} = 0.66$] and nature vs. indoor [b = 1.04, SE = 0.19, t = 5.58, p < 0.001, $d_{rm} = 1.05$] but not urban vs. indoor [b = 0.35, SE = 0.17, t = 2.03, p = 0.33, $d_{rm} = 0.34$] was associated with higher ratings of distraction away from the painful episode. The same pattern was found for ratings of tolerance towards the painful stimulus, indicating significantly higher ratings of tolerance in nature when comparing nature vs. urban [b = 1.10, SE = 0.20, t = 5.41, p < 0.001, $d_{rm} = 1.02$] and nature vs. indoor [b = 1.32, SE = 0.19, t = 7.12, p < 0.001, $d_{rm} = 1.33$] but not urban vs. indoor [b = 0.22, SE = 0.17, t = 1.32, p = 0.77, $d_{rm} = 0.25$]. The results mirror the findings observed using immediate self-reported pain by revealing effects specific to the nature condition. Compared to immediate ratings, they indicate effect sizes in the medium to high range when comparing nature to the other two conditions.

**Neural Responses to Pain.** To establish whether our experimental design led to activity differences in regions and multivariate signature responses associated with pain processing, irrespective of (i.e., statistically orthogonal to) the conditions of interest, we performed a whole-brain, a region of interest (ROI), and a signature response analysis. For all analyses, a pain>no-pain contrast was created and thresholded using familywise-error (FWE) correction at voxel-level (p < .05). The contrast was calculated across all three environments, thus encompassing a total of 48 painful and 48 non-painful shocks. First, conducting the whole-brain analysis revealed extensive hemodynamic activity across several brain areas (see Table S3 and Figure S1), including, among others, the anterior and posterior insula (aINS & pINS; bilateral), right primary somatosensory cortex (S1), secondary somatosensory cortex (S2; bilateral), anterior (ACC), and middle cingulate gyrus (MCC), superior frontal gyrus (SFG; including the supplementary motor area), cerebellum (bilateral), superior parietal lobe (SPL; bilateral), thalamus (bilateral) and periaqueductal grey (PAG). Second, we calculated a ROI analysis using preregistered sphere-based ROIs (for details see *Materials and Methods*) extracted from previous meta-analyses on acute pain and studies using a similar pain paradigm conducted in our research group (for references see main manuscript). These were the same ROIs as used in the main analyses of the manuscript, i.e., amygdala, anterior midcingulate cortex (aMCC), anterior (aINS) and posterior insula (pINS), medial prefrontal cortex (mPFC), primary (S1) and secondary (S2), periaqueductal grey (PAG), superior parietal lobe (SPL), and thalamus. We extracted the percent signal change of each ROI using the MarsBar toolbox (see reference 46 in main text) for the pain>no-pain contrast and ran separate dependent t-tests comparing differences in signal change against zero (see Table S4). The p-values for the ROI analyses were corrected by the overall number of investigated ROIs (p = .05/10 = .005) to account for multiple comparisons (p-values in Table S4 represent adjusted values). As can be seen in Table S4, all ROIs except the left primary somatosensory cortex (ipsilateral to the painful stimulation) revealed a significant response. Third, we calculated the signature response for the NPS and SIIPS1 using the dot product of the contrast image (pain>no-pain) with the pattern map of the NPS or SIIPS1. Again, dependent t-tests were calculated comparing the signature response against zero, which revealed a significant result for the NPS ($t_{(48)}$ = 9.23, $p$ < .001) and SIIPS1 ($t_{(48)}$ = 4.53, $p$ < .001). Collectively, our analyses validate the efficacy of our pain paradigm in activating brain regions and multivariate signature responses typically associated with the first-hand experience of pain.

After establishing that our paradigm effectively led to neural signal changes associated with the first-hand experience of pain, we conducted our main analyses. The effects of interest are reported in the main manuscript. Here, we provide details regarding the remaining effects and detailed information of each LMM conducted.

First, we calculated a LMM using the signature responses (NPS and SIIPS1) as a dependent variable (see Table S5), to be predicted by the fixed effect of environment (nature as a reference), signature (NPS as reference), and their interaction (with random slopes and intercepts for environment and signature by participant). The signature response of the NPS and the SIIPS1 was standardized prior to calculating the model, resulting in a non-significant main effect of environment [$F_{(2,48)}$ = 1.25, p = 0.296)] and signature [$F_{(1,48)}$ = 0.00, p = 1.00)]; note that the second main effect cannot be meaningfully interpreted, as the responses were standardized in advance. However, since we were not interested in whether the overall response of both signatures, irrespective of the environment, would differ, we deliberately specified the model in this way. Importantly, and as mentioned in the main manuscript, there was a significant interaction effect between environment and signature [$F_{(2,96)}$ = 6.04, p = 0.003)]. Planned pairwise comparisons revealed that there was a significant decrease in the NPS response during nature compared to the urban or indoor condition (see main manuscript). For the SIIPS1, no significant effects for the nature vs. urban [β = 0.24, SE = 0.16, t = 1.46, p = 0.926 one-tailed, $d_{rm}$ = .25] or indoor [β = -0.17, SE = 0.18, t = -0.94, p = 0.174 one-tailed, $d_{rm}$ = -.16] comparison were found, but a significant difference when comparing urban vs. indoor emerged [β = -0.41, SE = 0.16, t = 2.49, p = 0.014, $d_{rm}$ = -.41].

As indicated above, we calculated a second LMM using a different operationalization of lower-level vs. higher-level pain components. For this, we specified a preregistered LMM in which, instead of the NPS and SIIPS1, all ROI responses related to sensory-discriminative (i.e., S1, S2, pINS, SPL) or affective-motivational (i.e., aMCC, aINS, mPFC, dlPFC, PAG) components were used as the dependent variable simultaneously. We included environment (nature as a reference), component (sensory-discriminative as a reference), and their interaction as fixed effects. Using this model (see details in Table S6), we found a trend results for the main effect of environment on the overall pooled ROI response [$F_{(2,48)}$ = 2.84, p = 0.068], a significant result for the main effect of component [$F_{(1,48)}$ = 14.48, p = 0.003], and importantly and as expected a trend interaction effect of environment*component [$F_{(2,96)}$ = 2.72, p = 0.070]. Using planned pairwise contrasts revealed that there was a significant difference when comparing nature vs. urban [b = -0.50, SE = 0.18, t = -2.81, p = 0.006, $d_{rm}$ = -0.49] and nature vs. indoor [b = -0.45, SE = 0.21, t = -2.14, p = 0.036, $d_{rm}$ = -0.36] but not urban vs. indoor [b = 0.05, SE = 0.21, t = 0.26, p = 0.79, $d_{rm}$ = 0.07] when investigating the pooled activity of sensory-discriminative ROIs. For pooled activity of affective-motivational ROIs, no significant difference between nature vs. urban [b = -0.26, SE = 0.18, t = -1.43, p = 0.15, $d_{rm}$ = -0.19], nature vs. indoor [b = -0.19, SE = 0.21, t = -0.93, p = 0.35, $d_{rm}$ = -0.19] or urban vs. indoor emerged [b = 0.06, SE = 0.21, t = 0.29, p = 0.77, $d_{rm}$ = -0.01].

Importantly, these results mirror the analysis presented in the main manuscript by showing that it is the lower-level sensory-discriminative components rather than the higher-level cognitive-emotional components that nature stimuli act on. Thus, using the initially planned analysis, we further corroborate the findings of the main manuscript by showing that nature also reduces pain when using a different indicator of lower-level pain processing.

Second, we ran an individual LMM for each of the preregistered ROIs. For ROIs with a sphere covering voxels in both hemispheres, we ran a LMM using the ROI response as the dependent variable to be predicted by the fixed effect of environment (nature as a reference, with random intercepts for participants). For ROIs with a separate sphere in each hemisphere, we used the ROI response of both hemispheres as a dependent variable to be predicted by the fixed effect of environment (nature as a reference), hemisphere (left as reference), and their interaction (with random slopes and intercepts for environment and hemisphere by participant).

For the main effect of environment we observed the following results for our ROIs: amygdala [$F_{(2,48)}$ = 2.68, p = 0.078)], aINS [$F_{(2,48)}$ = 2.39, p = 0.102)], aMCC [$F_{(2,96)}$ = 0.59, p = 0.555)], mPFC [$F_{(2,96)}$ = 0.28, p = 0.756)], PAG [$F_{(2,96)}$ = 0.85, p = 0.431)], pINS [$F_{(2,48)}$ = 9.28, p < 0.001)], S1 [$F_{(2,48)}$ = 0.97, p = 0.386)], S2 [$F_{(2,48)}$ = 5.16, p = 0.009)], SPL [$F_{(2,48)}$ = 2.07, p = 0.136)], thalamus [$F_{(2,48)}$ = 5.54, p = 0.006)]. Planned pairwise comparisons contrasting nature vs. urban and nature vs. indoor conditions for the ROIs with significant or trend level main effects of environment (i.e., amygdala, pINS, S2, thalamus) are presented in the main manuscript. Planned pairwise contrasts revealed no significant differences when comparing urban vs. indoor in these ROIs indicating that this effect was specific for comparisons involving the nature condition: Thalamus [b = -0.10, SE = 0.11, t = -0.95, p = 0.346, $d_{rm}$ = -0.13], S2 [b = 0.11, SE = 0.21, t = 0.51, p = 0.615, $d_{rm}$ = 0.08], pINS [b = 0.58, SE = 0.25, t = 2.29, p = .105 $d_{rm}$ = 0.41] and the amygdala [b = 0.05, SE = 0.09, t = 0.55, p = 0.615, $d_{rm}$ = 0.10]. For the ROIs with two spheres the main effect of hemisphere revealed the following results: amygdala [$F_{(1,68.22)}$ = 0.18, p = 0.676)], aINS [$F_{(1,47.99)}$ = 1.02, p = 0.317)], pINS [$F_{(1,78.84)}$ = 5.02, p = 0.027)], S1 [$F_{(1,48)}$ = 54.25, p < 0.001)], S2 [$F_{(1,48)}$ = 11.52, p = 0.001)], SPL [$F_{(1,48)}$ = 21.47, p < 0.001)], thalamus [$F_{(1,48)}$ = 5.19, p = 0.027)]. Regarding the same set of ROIs the interaction effect of environment*hemisphere revealed the following results: amygdala [$F_{(2,143.99)}$ = 3.42, p = 0.035)], aINS [$F_{(2,96)}$ = 1.17, p = 0.313)], pINS [$F_{(2,144)}$ = 1.93, p = 0.148)], S1 [$F_{(2,96)}$ = 1.70, p = 0.187)], S2 [$F_{(2,95.99)}$ = 5.53, p = 0.005)], SPL [$F_{(2,96)}$ = 0.21, p = 0.809)], thalamus [F(2,96) = 1.59, p = 0.208)]. Thus, concerning differences in the hemispheres, we found significant interactions of environment*hemisphere in the S2 and amygdala. Post hoc tests indicated that differences in these ROIs were found in the left hemisphere (i.e., ipsilateral to the stimulated hand). All model details separated by ROI can be found in Table S7 – Table S16.

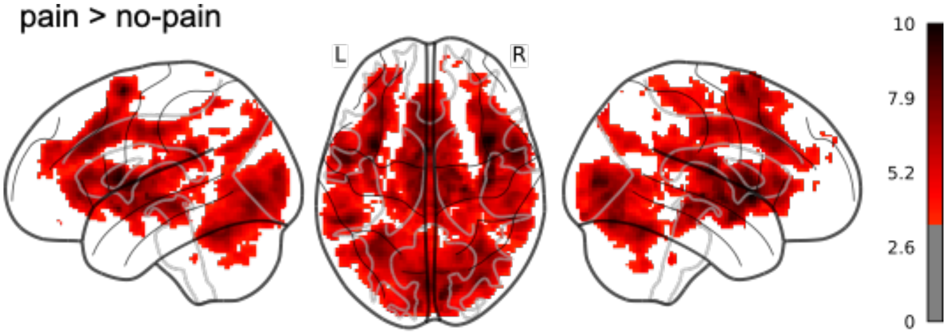


**Fig. S1.** Whole-brain analysis using the pain>no-pain contrast across all three environments. Note that due to analyses across all three environments, a comparably large number of 48 painful and 48 non-painful shocks were contrasted against each other. The glass brain shows activity changes in the hemodynamic response triggered by electrical stimulations of the left hand. We observed increased activity in a wide variety of brain regions associated with processing first-hand pain. A whole-brain display of the statistical activation map is shown, FWE-corrected at $p < .05$; L = left hemisphere, R = right hemisphere; Color bar depicts t-values.

**Table S1.** Linear mixed model results for immediate subjective pain ratings (i.e., intensity and unpleasantness).

| Fixed Effects | | | | | |
|---|---|---|---|---|---|
| | Estimate | SE | 95% CI | t | p |
| Intercept | 5.47 | 0.14 | 5.202 – 5.736 | 40.38 | <0.001 |
| Nat vs. Urb | 0.25 | 0.12 | 0.019 – 0.476 | 2.14 | 0.037 |
| Nat vs. Ind | 0.29 | 0.11 | 0.077 – 0.509 | 2.67 | 0.009 |
| Int vs. Unp | -0.81 | 0.14 | -1.081 – -0.538 | -5.87 | <0.001 |
| Nat vs. Urb * Int vs. Unp | 0.58 | 0.14 | 0.311 – 0.851 | 4.23 | <0.001 |
| Nat vs. Ind * Int vs. Unp | 0.37 | 0.13 | 0.109 – 0.623 | 2.81 | 0.006 |

| Random Effects | | | | | | | |
|---|---|---|---|---|---|---|---|
| | Variance | S.D. | Correlation | | | | |
| Participant (Intercept) | 0.732 | 0.855 | | | | | |
| Nat vs. Urb (Slope) | 0.326 | 0.571 | -.33 | | | | |
| Nat vs. Ind (Slope) | 0.255 | 0.505 | -.24 | .39 | | | |
| Int vs. Unp (Slope) | 0.597 | 0.772 | .20 | -.45 | -.16 | | |
| Nat vs. Urb (Slope) * Int vs. Unp | 0.258 | 0.508 | -.31 | .55 | .27 | -.28 | |
| Nat vs. Ind (Slope) * Int vs. Unp | 0.169 | 0.411 | -.16 | .59 | .89 | -.14 | .62 |

| Model fit | | |
|---|---|---|
| $R^2$ | Marginal | Conditional |
| | 0.12 | 0.55 |

*Note*: p-values for fixed effects calculated using Satterthwaites approximations.
Confidence Intervals have been calculated using the Wald method.
Model equation: Immediate_Pain ~ Environment*Content + (1+Environment*Content|Participant)
Nat = Nature, Urb = Urban, Ind = Indoor, Int = Intensity, Unp = Unpleasant.

**Table S2.** Linear mixed model results for retrospective subjective pain ratings (i.e., distraction and tolerance).

| Fixed Effects | | | | | |
|---|---|---|---|---|---|
| | Estimate | SE | 95% CI | t | p |
| Intercept | 3.02 | 0.57 | 1.89 – 4.15 | 5.27 | <0.001 |
| Nat vs. Urb | 0.94 | 0.75 | -0.54 – 2.41 | 1.25 | 0.212 |
| Nat vs. Ind | 0.10 | 0.74 | -1.36 – 1.57 | 0.14 | 0.891 |
| Dis vs. Tol | ~0.00 | 0.12 | -0.24 – 0.24 | 0.00 | 1.000 |
| Nat vs. Urb * Dis vs. Tol | -0.41 | 0.16 | -0.73 – -0.09 | -2.53 | 0.013 |
| Nat vs. Ind * Dis vs. Tol | -0.29 | 0.16 | -0.60 – 0.03 | -1.77 | 0.079 |

| Random Effects | | | | | |
|---|---|---|---|---|---|
| | Variance | S.D. | Correlation | | |
| Participant (Intercept) | 3.087 | 1.757 | | | |
| Nat vs. Urb (Slope) | 1.400 | 1.183 | -.29 | | |
| Nat vs. Ind (Slope) | 1.064 | 1.031 | -.41 | .69 | |
| Dis vs. Tol (Slope) | 0.099 | 0.315 | -.85 | -.11 | -.01 |

| Model fit | | |
|---|---|---|
| $R^2$ | Marginal | Conditional |
| | 0.22 | 0.75 |

*Note*: p-values for fixed effects calculated using Satterthwaites approximations. Confidence Intervals have been calculated using the Wald method.
Model equation: Retrospective_Pain ~ Environment*Content + (1+Environment+Content|Participant).
Nat = Nature, Urb = Urban, Ind = Indoor, Dis = Distraction, Tol = Tolerance.

**Table S3.** Whole-brain results for the contrast pain>no-pain across all environments.

| Cluster and peak levels | h | *x* | *y* | *z* | *t*-value | *p*-value |
|---|---|---|---|---|---|---|
| **Cluster 1** ($k$ = 62538) | | | | | | < .001 |
| Insular cortex | R | 38 | 10 | -2 | 10.49 | < .001 |
| Insular Cortex | L | -34 | 8 | 8 | 10.23 | < .001 |
| Superior Frontal Gyrus | L | -2 | 4 | 64 | 10.17 | < .001 |

*Note:* Significant clusters resulting from the contrast [*pain>no-pain*]. h = hemisphere, $k$ = cluster size, MNI coordinates x, y, z, *t*-value and *p*-value (FWE-corrected at $p < .05$, cluster level, $k >$ 213). Peak coordinates were labelled according to the automated anatomical labelling (AAL2) atlas (11).

**Table S4.** Region of interest results for the contrast pain>no-pain.

| **Region** | df | *t*-value | *p*-value |
|---|---|---|---|
| Amygdala (left) | 48 | 5.70 | < .001 |
| Amygdala (right) | 48 | 4.62 | < .001 |
| Anterior insula (left) | 48 | 8.36 | < .001 |
| Anterior insula (right) | 48 | 8.38 | < .001 |
| Anterior midcingulate cortex | 48 | 9.05 | < .001 |
| Medial prefrontal cortex | 48 | 4.72 | < .001 |
| Periaqueductal grey | 48 | 7.50 | < .001 |
| Posterior insula (left) | 48 | 7.59 | < .001 |
| Posterior insula (right) | 48 | 7.95 | < .001 |
| Primary somatosensory cortex (left) | 48 | 1.44 | .155 |
| Primary somatosensory cortex (right) | 48 | 3.63 | < .01 |
| Secondary somatosensory cortex (left) | 48 | 7.16 | < .001 |
| Secondary somatosensory cortex (right) | 48 | 9.15 | < .001 |
| Superior parietal lobe (left) | 48 | 5.01 | < .001 |
| Superior parietal lobe (right) | 48 | 6.89 | < .001 |
| Thalamus (left) | 48 | 6.87 | < .001 |
| Thalamus (right) | 48 | 7.39 | < .001 |

*Note:* All p-values represent adjusted values (Bonferroni corrected; $p = .005$)

**Table S5.** Linear mixed model results for multivariate signature responses (i.e., NPS and SIIPS1).

| Fixed Effects | | | | | |
|---|---|---|---|---|---|
| | Estimate | SE | 95% CI | t | p |
| Intercept | -0.22 | 0.13 | -0.47 – 0.03 | -1.77 | 0.081 |
| Nat vs. Urb | 0.37 | 0.16 | 0.05 – 0.69 | 2.26 | 0.025 |
| Nat vs. Ind | 0.30 | 0.18 | -0.05 – 0.66 | 1.67 | 0.099 |
| NPS vs. SIIPS1 | 0.24 | 0.14 | -0.03 – 0.53 | 1.73 | 0.085 |
| Nat vs. Urb * NPS vs. SIIPS1 | -0.61 | 0.18 | -0.97 – -0.25 | -3.30 | 0.001 |
| Nat vs. Ind * NPS vs. SIIPS1 | -0.13 | 0.18 | -0.49 – 0.23 | -0.72 | 0.474 |

| Random Effects | | | | | |
|---|---|---|---|---|---|
| | Variance | S.D. | Correlation | | |
| Participant (Intercept) | 0.371 | 0.609 | | | |
| Nat vs. Urb (Slope) | 0.469 | 0.685 | -.54 | | |
| Nat vs. Ind (Slope) | 0.795 | 0.891 | -.23 | .63 | |
| NPS vs. SIIPS1 (Slope) | 0.162 | 0.403 | -.11 | -.22 | -.16 |

| Model fit | | |
|---|---|---|
| $R^2$ | Marginal | Conditional |
| | 0.03 | 0.59 |

*Note*: p-values for fixed effects calculated using Satterthwaites approximations. Confidence Intervals have been calculated using the Wald method.
Model equation: Signature_Response ~ Environment*Signature + (1+Environment+Signature|Participant).
Nat = Nature, Urb = Urban, Ind = Indoor, NPS = neurologic pain signature, SIIPS1 = stimulus intensity independent pain signature-1.

**Table S6.** Linear mixed model results for pooled ROI responses (i.e., ROIs clustered into sensory-discriminative and affective-motivational components).

| Fixed Effects | | | | | |
|---|---|---|---|---|---|
| | Estimate | SE | 95% CI | t | p |
| Intercept | 1.16 | 0.18 | 0.80 – 1.53 | 6.53 | <0.001 |
| Nat vs. Urb | 0.26 | 0.18 | -0.14 – 0.58 | 1.53 | 0.156 |
| Nat vs. Ind | 0.19 | 0.20 | -0.21 – 0.60 | 0.93 | 0.353 |
| Aff-Mot vs. Sen-Dis | -0.43 | 0.11 | -0.72 – -0.30 | -3.86 | <0.001 |
| Nat vs. Urb * Aff-Mot vs. Sen-Dis | 0.24 | 0.12 | 0.08 – 0.50 | 1.99 | 0.049 |
| Nat vs. Ind * Aff-Mot vs. Sen-Dis | 0.25 | 0.12 | 0.05 – 0.46 | 2.05 | 0.043 |

| Random Effects | | | | | |
|---|---|---|---|---|---|
| | Variance | S.D. | Correlation | | |
| Participant (Intercept) | 1.355 | 1.164 | | | |
| Nat vs. Urb (Slope) | 1.192 | 1.092 | -.39 | | |
| Nat vs. Ind (Slope) | 1.758 | 1.326 | -.38 | .39 | |
| Aff-Mot vs. Sen-Dis (Slope) | 0.244 | 0.494 | -.51 | .00 | .26 |

| Model fit | | |
|---|---|---|
| $R^2$ | Marginal | Conditional |
| | 0.03 | 0.89 |

*Note*: p-values for fixed effects calculated using Satterthwaites approximations. Confidence Intervals have been calculated using the Wald method.
Model equation: Pooled_ROI_Response ~ Environment*Component + (1+Environment+Component|Participant).
Nat = Nature, Urb = Urban, Ind = Indoor, Aff-Mot = affective-motivational, Sen-Dis = sensory-discriminative.

**Table S7.** Linear mixed model results for the amygdala response.

| Fixed Effects | | | | | |
|---|---|---|---|---|---|
| | Estimate | SE | 95% CI | t | p |
| Intercept | 0.27 | 0.06 | 0.16 – 0.39 | 4.58 | <0.001 |
| Nat vs. Urb | 0.25 | 0.09 | 0.06 – 0.44 | 2.58 | 0.012 |
| Nat vs. Ind | 0.11 | 0.07 | -0.03 – 0.26 | 1.55 | 0.124 |
| Left vs. Right | 0.04 | 0.05 | -0.07 – 0.14 | 0.68 | 0.494 |
| Nat vs. Urb * Left vs. Right | -0.16 | 0.07 | -0.30 – -0.02 | -2.23 | 0.027 |
| Nat vs. Ind * Left vs. Right | 0.00 | 0.07 | -0.13 – 0.14 | 0.07 | 0.945 |

| Random Effects | | | | | |
|---|---|---|---|---|---|
| | Variance | S.D. | Correlation | | |
| Participant (Intercept) | 0.115 | 0.338 | | | |
| Nat vs. Urb (Slope) | 0.343 | 0.585 | -.21 | | |
| Nat vs. Ind (Slope) | 0.139 | 0.373 | -.37 | .22 | |
| Left vs. Right (Slope) | 0.017 | 0.131 | -.21 | -.75 | .45 |

| Model fit | | |
|---|---|---|
| $R^2$ | Marginal | Conditional |
| | 0.03 | 0.77 |

*Note*: p-values for fixed effects calculated using Satterthwaites approximations. Confidence Intervals have been calculated using the Wald method.
Model equation: Amy_Response ~ Environment*Hemisphere + (1+Environment+Hemisphere|Participant).
Nat = Nature, Urb = Urban, Ind = Indoor.

**Table S8.** Linear mixed model results for the anterior insula (aINS) response.

| Fixed Effects | | | | | |
|---|---|---|---|---|---|
| | Estimate | SE | 95% CI | t | p |
| Intercept | 0.92 | 0.16 | 0.61 – 1.23 | 5.88 | <0.001 |
| Nat vs. Urb | 0.22 | 0.18 | -0.13 – 0.57 | 1.24 | 0.216 |
| Nat vs. Ind | 0.43 | 0.19 | 0.05 – 0.81 | 2.22 | 0.031 |
| Left vs. Right | 0.05 | 0.07 | -0.09 – 0.20 | 0.78 | 0.436 |
| Nat vs. Urb * Left vs. Right | 0.06 | 0.08 | -0.10 – 0.22 | 0.73 | 0.466 |
| Nat vs. Ind * Left vs. Right | -0.07 | 0.08 | -0.23 – 0.10 | -0.80 | 0.423 |

| Random Effects | | | | | |
|---|---|---|---|---|---|
| | Variance | S.D. | Correlation | | |
| Participant (Intercept) | 1.115 | 1.056 | | | |
| Nat vs. Urb (Slope) | 1.398 | 1.182 | -.53 | | |
| Nat vs. Ind (Slope) | 1.680 | 1.296 | -.33 | .41 | |
| Left vs. Right (Slope) | 0.086 | 0.293 | -.16 | .12 | -.03 |

| Model fit | | |
|---|---|---|
| $R^2$ | Marginal | Conditional |
| | 0.02 | 0.94 |

*Note*: p-values for fixed effects calculated using Satterthwaites approximations. Confidence Intervals have been calculated using the Wald method.
Model equation: aINS_Response ~ Environment*Hemisphere + (1+Environment+Hemisphere|Participant).
Nat = Nature, Urb = Urban, Ind = Indoor.

**Table S9.** Linear mixed model results for the anterior midcingulate cortex (aMCC) response.

| Fixed Effects | | | | | |
|---|---|---|---|---|---|
| | Estimate | SE | 95% CI | t | p |
| Intercept | 1.99 | 0.29 | 1.41 – 2.58 | 6.72 | <0.001 |
| Nat vs. Urb | 0.19 | 0.31 | -0.41 – 0.80 | 0.64 | 0.521 |
| Nat vs. Ind | 0.33 | 0.31 | -0.27 – 0.94 | 1.08 | 0.282 |

| Random Effects | | |
|---|---|---|
| | Variance | S.D. |
| Participant (Intercept) | 2.019 | 1.42 |

| Model fit | | |
|---|---|---|
| $R^2$ | Marginal | Conditional |
| | 0.004 | 0.47 |

*Note*: p-values for fixed effects calculated using Satterthwaites approximations. Confidence Intervals have been calculated using the Wald method.
Model equation: aMCC_Response ~ Environment + (1+Environment |Participant).
Nat = Nature, Urb = Urban, Ind = Indoor.

**Table S10.** Linear mixed model results for the medial prefrontal cortex (mPFC) response.

| Fixed Effects | | | | | |
|---|---|---|---|---|---|
| | Estimate | SE | 95% CI | t | p |
| Intercept | 0.98 | 0.22 | 0.50 – 1.46 | 4.35 | <0.001 |
| Nat vs. Urb | 0.15 | 0.27 | -0.33 – 0.85 | 0.57 | 0.569 |
| Nat vs. Ind | -0.03 | 0.27 | -0.55 – 0.62 | -0.13 | 0.896 |

| Random Effects | | |
|---|---|---|
| | Variance | S.D. |
| Participant (Intercept) | 0.651 | 0.81 |

| Model fit | | |
|---|---|---|
| $R^2$ | Marginal | Conditional |
| | 0.005 | 0.25 |

*Note*: p-values for fixed effects calculated using Satterthwaites approximations. Confidence Intervals have been calculated using the Wald method.
Model equation: mPFC_Response ~ Environment + (1+Environment|Participant).
Nat = Nature, Urb = Urban, Ind = Indoor.

**Table S11.** Linear mixed model results for the periaqueductal grey (PAG) response.

| Fixed Effects | | | | | |
|---|---|---|---|---|---|
| | Estimate | SE | 95% CI | t | p |
| Intercept | 0.54 | 0.11 | 0.32 – 0.77 | 4.84 | <0.001 |
| Nat vs. Urb | 0.18 | 0.13 | -0.09 – 0.45 | 1.29 | 0.197 |
| Nat vs. Ind | 0.07 | 0.13 | -0.20 – 0.35 | 0.54 | 0.591 |

| Random Effects | | |
|---|---|---|
| | Variance | S.D. |
| Participant (Intercept) | 0.152 | 0.39 |

| Model fit | | |
|---|---|---|
| $R^2$ | Marginal | Conditional |
| | 0.009 | 0.25 |

*Note*: p-values for fixed effects calculated using Satterthwaites approximations. Confidence Intervals have been calculated using the Wald method.
Model equation: PAG_Response ~ Environment + (1+Environment|Participant).
Nat = Nature, Urb = Urban, Ind = Indoor.

**Table S12.** Linear mixed model results for the posterior insula (pINS) response.

| Fixed Effects | | | | | |
|---|---|---|---|---|---|
| | Estimate | SE | 95% CI | t | p |
| Intercept | 0.46 | 0.14 | 0.18 – 0.74 | 3.26 | 0.001 |
| Nat vs. Urb | 1.10 | 0.23 | 0.64 – 1.57 | 4.65 | <0.001 |
| Nat vs. Ind | 0.45 | 0.21 | 0.04 – 0.87 | 2.14 | 0.036 |
| Left vs. Right | 0.29 | 0.11 | 0.08 – 0.50 | 2.73 | 0.007 |
| Nat vs. Urb * Left vs. Right | -0.28 | 0.14 | -0.56 - 0.00 | -1.97 | 0.051 |
| Nat vs. Ind * Left vs. Right | -0.14 | 0.14 | -0.42 – 0.14 | -0.98 | 0.327 |

| Random Effects | | | | | |
|---|---|---|---|---|---|
| | Variance | S.D. | Correlation | | |
| Participant (Intercept) | 0.727 | 0.853 | | | |
| Nat vs. Urb (Slope) | 2.265 | 1.505 | -.37 | | |
| Nat vs. Ind (Slope) | 1.682 | 1.297 | -.37 | .26 | |
| Left vs. Right (Slope) | 0.046 | 0.216 | .56 | -.69 | .31 |

| Model fit | | |
|---|---|---|
| $R^2$ | Marginal | Conditional |
| | 0.09 | 0.87 |

*Note*: p-values for fixed effects calculated using Satterthwaites approximations. Confidence Intervals have been calculated using the Wald method.
Model equation: pINS_Response ~ Environment*Hemisphere + (1+Environment+Hemisphere|Participant).
Nat = Nature, Urb = Urban, Ind = Indoor.

**Table S13.** Linear mixed model results for the primary somatosensory cortex (S1) response.

| Fixed Effects | | | | | |
|---|---|---|---|---|---|
| | Estimate | SE | 95% CI | t | p |
| Intercept | 0.06 | 0.16 | -0.26 – 0.38 | 0.35 | 0.728 |
| Nat vs. Urb | 0.27 | 0.19 | -0.10 – 0.64 | 1.44 | 0.154 |
| Nat vs. Ind | 0.15 | 0.24 | -0.32 – 0.62 | 0.62 | 0.535 |
| Left vs. Right | 0.69 | 0.15 | 0.40 – 0.98 | 4.67 | <0.001 |
| Nat vs. Urb * Left vs. Right | -0.16 | 0.19 | -0.54 – 0.23 | -0.81 | 0.420 |
| Nat vs. Ind * Left vs. Right | 0.20 | 0.19 | -0.18 – 0.58 | 1.03 | 0.305 |

| Random Effects | | | | | |
|---|---|---|---|---|---|
| | Variance | S.D. | Correlation | | |
| Participant (Intercept) | 0.833 | 0.912 | | | |
| Nat vs. Urb (Slope) | 0.822 | 0.907 | -.39 | | |
| Nat vs. Ind (Slope) | 1.871 | 1.367 | -.43 | .38 | |
| Left vs. Right (Slope) | 0.137 | 0.369 | -.37 | .63 | .32 |

| Model fit | | |
|---|---|---|
| $R^2$ | Marginal | Conditional |
| | 0.08 | 0.75 |

*Note*: p-values for fixed effects calculated using Satterthwaites approximations. Confidence Intervals have been calculated using the Wald method.
Model equation: S1_Response ~ Environment*Hemisphere + (1+Environment+Hemisphere|Participant).
Nat = Nature, Urb = Urban, Ind = Indoor.

**Table S14.** Linear mixed model results for the secondary somatosensory cortex (S2) response.

| Fixed Effects | | | | | |
|---|---|---|---|---|---|
| | Estimate | SE | 95% CI | t | p |
| Intercept | 0.53 | 0.14 | 0.25 – 0.80 | 3.71 | <0.001 |
| Nat vs. Urb | 0.63 | 0.18 | 0.26 – 0.99 | 3.40 | 0.001 |
| Nat vs. Ind | 0.49 | 0.17 | 0.15 – 0.84 | 2.84 | 0.005 |
| Left vs. Right | 0.33 | 0.07 | 0.19 – 0.47 | 4.67 | <0.001 |
| Nat vs. Urb * Left vs. Right | -0.25 | 0.08 | -0.41 – -0.09 | -3.15 | 0.002 |
| Nat vs. Ind * Left vs. Right | -0.20 | 0.08 | -0.36 – -0.04 | -2.49 | 0.014 |

| Random Effects | | | | | |
|---|---|---|---|---|---|
| | Variance | S.D. | Correlation | | |
| Participant (Intercept) | 0.902 | 0.950 | | | |
| Nat vs. Urb (Slope) | 1.497 | 1.223 | -.47 | | |
| Nat vs. Ind (Slope) | 1.320 | 1.149 | -.37 | .27 | |
| Left vs. Right (Slope) | 0.088 | 0.298 | -.33 | .20 | .11 |

| Model fit | | |
|---|---|---|
| $R^2$ | Marginal | Conditional |
| | 0.04 | 0.94 |

*Note*: p-values for fixed effects calculated using Satterthwaites approximations. Confidence Intervals have been calculated using the Wald method.
Model equation: S2_Response ~ Environment*Hemisphere + (1+Environment+Hemisphere|Participant).
Nat = Nature, Urb = Urban, Ind = Indoor.

**Table S15.** Linear mixed model results for the superior parietal lobe (SPL) response.

| Fixed Effects | | | | | |
|---|---|---|---|---|---|
| | Estimate | SE | 95% CI | t | p |
| Intercept | 0.77 | 0.25 | 0.29 – 1.30 | 3.11 | 0.002 |
| Nat vs. Urb | 0.36 | 0.31 | -0.20 – 1.01 | 1.18 | 0.244 |
| Nat vs. Ind | 0.69 | 0.39 | -0.03 – 1.51 | 1.80 | 0.077 |
| Left vs. Right | 0.82 | 0.23 | 0.36 – 1.26 | 3.62 | <0.001 |
| Nat vs. Urb * Left vs. Right | -0.06 | 0.24 | -0.60 – 0.38 | -0.28 | 0.778 |
| Nat vs. Ind * Left vs. Right | 0.09 | 0.24 | -0.45 – 0.53 | 0.36 | 0.715 |

| Random Effects | | | | | |
|---|---|---|---|---|---|
| | Variance | S.D. | Correlation | | |
| Participant (Intercept) | 2.328 | 1.526 | | | |
| Nat vs. Urb (Slope) | 3.172 | 1.781 | -.60 | | |
| Nat vs. Ind (Slope) | 5.830 | 2.413 | -.44 | .45 | |
| Left vs. Right (Slope) | 1.104 | 1.051 | .27 | -.15 | .17 |

| Model fit | | |
|---|---|---|
| $R^2$ | Marginal | Conditional |
| | 0.05 | 0.86 |

*Note*: p-values for fixed effects calculated using Satterthwaites approximations. Confidence Intervals have been calculated using the Wald method.
Model equation: SPL_Response ~ Environment*Hemisphere + (1+Environment+Hemisphere|Participant).
Nat = Nature, Urb = Urban, Ind = Indoor.

**Table S16.** Linear mixed model results for the thalamus response.

| Fixed Effects | | | | | |
|---|---|---|---|---|---|
| | Estimate | SE | 95% CI | t | p |
| Intercept | 0.35 | 0.10 | 0.15 – 0.56 | 3.43 | 0.001 |
| Nat vs. Urb | 0.31 | 0.11 | 0.09 – 0.54 | 2.76 | 0.006 |
| Nat vs. Ind | 0.44 | 0.12 | 0.19 – 0.68 | 3.52 | <0.001 |
| Left vs. Right | 0.14 | 0.05 | 0.04 – 0.25 | 2.80 | 0.005 |
| Nat vs. Urb * Left vs. Right | -0.07 | 0.06 | -0.19 – 0.05 | -1.14 | 0.256 |
| Nat vs. Ind * Left vs. Right | -0.11 | 0.06 | -0.23 – 0.01 | -1.76 | 0.080 |

| Random Effects | | | | | |
|---|---|---|---|---|---|
| | Variance | S.D. | Correlation | | |
| Participant (Intercept) | 0.469 | 0.685 | | | |
| Nat vs. Urb (Slope) | 0.541 | 0.730 | -.54 | | |
| Nat vs. Ind (Slope) | 0.663 | 0.814 | -.43 | .55 | |
| Left vs. Right (Slope) | 0.036 | 0.191 | -.09 | .22 | .32 |

| Model fit | | |
|---|---|---|
| $R^2$ | Marginal | Conditional |
| | 0.05 | 0.93 |

*Note*: p-values for fixed effects calculated using Satterthwaites approximations. Confidence Intervals have been calculated using the Wald method.
Model equation: Thal_Response ~ Environment*Hemisphere + (1+Environment+Hemisphere|Participant).
Nat = Nature, Urb = Urban, Ind = Indoor.

**SI References**


1. F. Faul, E. Erdfelder, A. Buchner, A.-G. Lang, Statistical power analyses using G*Power 3.1: Tests for correlation and regression analyses. *Behav Res Methods* **41**, 1149–1160 (2009).
2. G. Chen, N. E. Adleman, Z. S. Saad, E. Leibenluft, R. W. Cox, Applications of multivariate modeling to neuroimaging group analysis: A comprehensive alternative to univariate general linear model. *Neuroimage* **99**, 571–588 (2014).
3. L. Meteyard, R. A. I. Davies, Best practice guidance for linear mixed-effects models in psychological science. *J Mem Lang* **112**, 104092 (2020).
4. M. Farzaneh, *et al.*, Comparative effect of nature-based sounds intervention and headphones intervention on pain severity after cesarean section: A prospective double-blind randomized trial. *Anesth Pain Med* **9**, e67835 (2019).
5. N. Lechtzin, *et al.*, A randomized trial of nature scenery and sounds versus urban scenery and sounds to reduce pain in adults undergoing bone marrow aspirate and biopsy. *J Altern Complement Med* **16**, 965–972 (2010).
6. D. Lee, *et al.*, Can visual distraction decrease the dose of patient-controlled sedation required during colonoscopy? A prospective randomized controlled trial. *Endoscopy* **36**, 197–201 (2004).
7. M. M. Y. Tse, J. K. F. Ng, J. W. Y. Chung, T. K. S. Wong, The effect of visual stimuli on pain threshold and tolerance. *J Clin Nurs* **11**, 462–469 (2002).
8. J. E. Letzen, *et al.*, Test-retest reliability of pain-related brain activity in healthy controls undergoing experimental thermal pain. *J Pain* **15**, 1008–1014 (2014).
9. R. L. Quiton, M. L. Keaser, J. Zhuo, R. P. Gullapalli, J. D. Greenspan, Intersession reliability of fMRI activation for heat pain and motor tasks. *Neuroimage Clin* **5**, 309–321 (2014).
10. J. Upadhyay, *et al.*, Test–retest reliability of evoked heat stimulation BOLD fMRI. *J Neurosci Methods* **253**, 38–46 (2015).
11. E. T. Rolls, M. Joliot, N. Tzourio-Mazoyer, Implementation of a new parcellation of the orbitofrontal cortex in the automated anatomical labeling atlas. *Neuroimage* **122**, 1–5 (2015).
